# The LEAFY floral regulator displays pioneer transcription factor properties

**DOI:** 10.1101/2021.01.07.425699

**Authors:** Xuelei Lai, Romain Blanc-Mathieu, Loïc GrandVuillemin, Ying Huang, Arnaud Stigliani, Jérémy Lucas, Emmanuel Thévenon, Jeanne Loue-Manifel, Hussein Daher, Eugenia Brun-Hernandez, Gilles Vachon, David Latrasse, Moussa Benhamed, Renaud Dumas, Chloe Zubieta, François Parcy

**Affiliations:** Laboratoire Physiologie Cellulaire et Végétale, Univ. Grenoble Alpes, CNRS, CEA, INRAE, IRIG-DBSCI-LPCV, 17 avenue des martyrs, F-38054, Grenoble, France; Université Paris-Saclay, CNRS, INRAE, Univ. Evry, Institute of Plant Sciences Paris-Saclay (IPS2), 91405, Orsay, France; The Bioinformatics Centre, Department of Biology and Biotech and Research Innovation Centre, University of Copenhagen, Ole Maaløes Vej 5, DK2200 Copenhagen N, Denmark; Laboratoire Reproduction et Développement des Plantes, Université de Lyon, ENS de Lyon, UCB Lyon 1, CNRS, INRA, F-69342 Lyon, France; Institut de Biologie Structurale, Université Grenoble Alpes, CEA, CNRS, Grenoble, France; Université de Paris, Institute of Plant Sciences Paris-Saclay (IPS2), F-75006 Paris, France

## Abstract

Pioneer transcription factors (TFs) are a special category of TFs with the capacity to bind to closed chromatin regions in which DNA is wrapped around histones and often highly methylated. Subsequently, they are able to modify the chromatin state to initiate gene expression. In plants, LEAFY (LFY) is a master floral regulator and has been suggested to act as a pioneer TF in Arabidopsis. Here, we demonstrate that LFY is able to bind both methylated and non-methylated DNA using a combination of *in vitro* genome-wide binding experiments and structural modeling. Comparisons between regions bound by LFY *in vivo* and chromatin accessibility data suggest that LFY binds a subset of regions occupied by nucleosomes. We confirm that LFY is able to bind nucleosomal DNA *in vitro* using reconstituted nucleosomes. Finally, we show that constitutive LFY expression in seedling tissues is sufficient to induce chromatin accessibility in the LFY direct target genes, *APETALA1* and *AGAMOUS*. Taken together, our study suggests that LFY possesses key pioneer TF features that contribute to launch the floral gene expression program.

## Introduction

Proper gene regulation is essential to all living organisms, controlling processes from basic development to environmental response. Gene regulation requires the finely orchestrated activity of transcription factors (TFs) that recognize specific DNA sequences in gene regulatory regions and activate or repress transcription of their target genes. While the binding of most TFs to DNA is restricted to accessible regions of the genome, a specific type of TF, called a “pioneer”, is able to access its cognate binding site even in closed, nucleosome-rich chromatin regions (Magnani et al., 2011; Iwafuchi-Doi and Zaret, 2014; Iwafuchi-Doi and Zaret, 2016; Zaret, 2020). The ability to bind nucleosomal DNA *in vivo* and *in vitro* is a defining characteristic of pioneer TFs and has been well-established for diverse mammalian pioneer TFs (Fernandez Garcia et al., 2019). As DNA in closed chromatin regions is often highly methylated, another emerging feature of pioneer TFs is their capability to bind DNA in a methylation insensitive manner (Zhu et al., 2016; Mayran and Drouin, 2018). Some pioneer TFs are even able to directly recruit DNA demethylases at methylated sites, thereby facilitating the remodeling of closed regions (Iwafuchi-Doi, 2018).

Pioneer TFs are often master regulators controlling developmental transitions, with the mammalian pluripotency factors OCT4, SOX2, and KLF4 representing some of the most well-studied (Soufi et al., 2015). These factors bind to closed chromatin regions and induce their opening or remodeling, so that genes they contain can be activated by the pioneer TFs themselves or by other TFs called settlers (Sherwood et al., 2014; Slattery et al., 2014). The modification of the chromatin landscape by pioneer TF can be accomplished either directly by triggering DNA detachment from nucleosomes (Dodonova et al., 2020; Michael et al., 2020), or indirectly by the recruitment of ATP-dependent cellular machineries, such as chromatin remodelers that remove or modify adjacent nucleosomes in order to prime downstream regulatory events (Hu et al., 2011; King and Klose, 2017). Such capacity to modify DNA accessibility is another defining feature of pioneer TFs (Iwafuchi-Doi and Zaret, 2014).

In plants, the only TF reported as pioneer TF so far is LEAFY COTYLEDON1 (LEC1), a seed specific TF involved in embryonic epigenetic reprogramming (Tao et al., 2017). LEC1 was shown to promote the initial establishment of an active chromatin state of its target gene in silenced chromatin and activate its expression de novo. Pioneer TF activity was also suggested for two types of factors controlling flower development, the MADS homeotic TFs (Pajoro et al., 2014; Denay et al., 2017) and the master floral regulator, LEAFY (LFY). The MADS TFs, including APETALA1 (AP1) and SEPALLATA3, were shown to be able to access closed chromatin regions to specify floral organs, and were thus postulated to act as pioneer TFs (Pajoro et al., 2014). However, mammalian MADS TFs do not seem to act as pioneer factors and thus the identification of AP1 and SEP3 as potential pioneers remains speculative (Sherwood et al., 2014). In contrast to the MADS TFs, one previous study suggest that LFY may have pioneer activity (Sayou et al., 2016). LFY is a master regulator specifying the floral identity of meristems. It directly induces the floral homeotic genes *AP1, APETALA3 (AP3)* and *AGAMOUS (AG)* (Parcy et al., 1998; Wagner, 1999; Lohmann et al., 2001; Chae et al., 2008; Yamaguchi et al., 2013; Chahtane et al., 2013). *AG* and *AP3* are known to be under the repression of Polycomb repressive complexes in seedlings (Goodrich et al., 1997; Turck et al., 2007; Calonje et al., 2008). This suggests that their activation during flower development requires modifications of their chromatin landscape and that the direct binding of LFY to their regulatory regions might trigger. Consistent with this, LFY was suggested to be able to access closed chromatin regions *in vivo* (Sayou et al., 2016). Moreover, LFY’s role is not confined to conferring a flower fate to meristems. It can also contribute to meristem emergence (Moyroud et al., 2010; Chahtane et al., 2013; Yamaguchi et al., 2013), and together with its co-regulators such as the homeodomain TF WUSCHEL or the F-Box protein UNUSUAL FLORAL ORGANS, it can even induce meristem formation from root or leaf tissue, respectively (Levin and Meyerowitz, 1995; Gallois et al., 2004; Risseeuw et al., 2013). Taken together, these data indicate that LFY has the full capability of reprogramming cell fate, a property often requiring pioneer activity. However, whether LFY is truly able to directly bind closed chromatin regions and change their status has yet to be demonstrated.

Here, we address the pioneer activity of LFY *in vitro* and *in vivo*. Firstly, we determined whether LFY binding was sensitive to DNA methylation. For this, we combined *in vitro* LFY genome-wide binding data using methylated and unmethylated genomic DNA and structural analysis. These experiments demonstrated that LFY binding is only mildly sensitive to DNA methylation. In order to test whether LFY binding was compatible with the presence of nucleosomes, we compared LFY binding data from chromatin immunoprecipitation sequencing (ChIP-seq) and chromatin accessibility data. Based on these comparisons, we found that LFY could access a number of closed chromatin regions and that LFY colocalizes with nucleosomes in some regions *in vivo*. Using electrophoretic mobility shift assays (EMSA), we further showed that LFY was able to directly bind nucleosomes *in vitro*. Finally, chromatin accessibility assays demonstrated that LFY constitutive expression was sufficient to increase chromatin accessibility in genomic regions including its known target genes *AP1* and *AG*. Taken together, these data establish that LFY is able to act as a pioneer TF in the regulation of important target genes critical for the establishment of floral fate.

## Results

### LFY is weakly sensitive to DNA methylation

In closed chromatin regions, DNA is packed within nucleosomes (McGinty and Tan, 2015) and its level of methylation is often higher than in open chromatin (Chodavarapu et al., 2010). Both the presence of nucleosome and DNA methylation usually reduce TFs access and their binding to target DNA (Yin et al., 2017; Klemm et al., 2019). In order to assess the effect of methylation on LFY binding to DNA, we applied DNA Affinity Purification sequencing (DAP-seq) (O’Malley et al., 2016). Similar to ChIP-seq, this technique allows the identification of the genomic regions bound by a TF but using naked DNA and a recombinant TF. We used Arabidopsis genomic DNA extracted from seedlings that was either PCR amplified (ampDAP, DNA cleared of methylation) or not amplified (DAP, DNA retaining methylation). Both experiments were performed in triplicates with high reproducibility (Supplemental Figure 1; Supplemental Table 1). As controls, we used two TFs described as methylation sensitive based on available ampDAP and DAP datasets (O’Malley et al., 2016) (Supplemental Figure 2). For each genomic region bound by a given TF, we plotted the DAP/ampDAP signal ratio as a function of the methylation density in the whole bound region (based on Arabidopsis seedling methylation maps (Zhang et al., 2016)). If the DNA binding of a TF is inhibited by methylation, we expect the DAP/ampDAP ratio to decrease when the methylation level increases. LFY DNA binding was much less affected by increasing methylation density than the two methylation sensitive TFs, (Figure 1A-C). To analyze more precisely the effect of methylation, we tested the correlation between the number of methylated cytosines within the best TF binding (TFBS) site, identified using position weight matrices in each bound region and the DAP/ampDAP ratio of bound regions. Whereas an increased number of methylated cytosines in TFBS strongly decreased the binding for the two methylation sensitive TFs in DAP relative to ampDAP, LFY binding was only mildly affected (Figure 1D-F). Finally, we designed a specific procedure to compute the effect of methylation on each individual cytosine possibly present in the best TFBS (Supplemental Figure 3–5). In the case of LFY, we identified two positions where the binding is increased by cytosine methylation (positions 4 on the forward DNA strand and 5 on the reverse), and other positions (2,3,7,8 on the forward strand and 1,3,4,9 on the reverse) where the binding is only mildly inhibited (Figure 1G). In contrast, methylation was inhibitory for the two methylation sensitive TFs in most positions where a cytosine can possibly be present (Figure 1H-I). Structural analysis of LFY DNA binding domain in complex with DNA (Hamès et al., 2008) provided a biochemical explanation of these positive and negative effects (Supplemental Figure 6). In particular, the hydrophobic contacts between LFY and DNA are likely to be enhanced by the presence of a methyl group in positions 4 and 5 of the LFY binding site (LFYBS), consistent with the DAP versus ampDAP analysis (Figure 1G).

**Figure 1:**
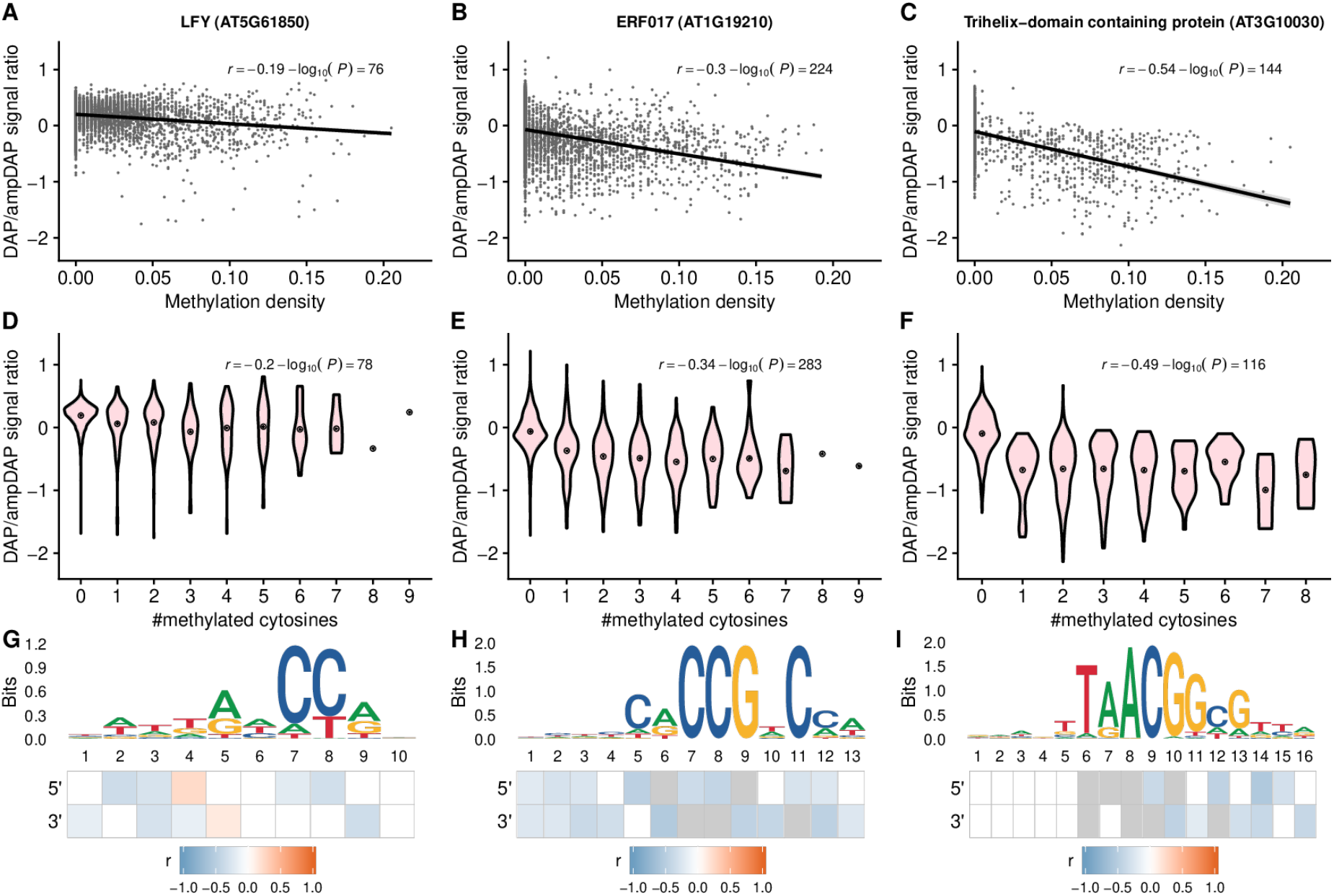
Cytosine methylation has a mild effect on LFY DNA-binding intensity. Effect of cytosine methylation on DNA binding for three transcription factors: LFY (left), ERF017 (middle) and a trihelix-domain containing protein (right). (A-C) Biplots between the DAP/ampDAP signal ratio (peak normalized read coverage in the DAP experiment divided to that in the ampDAP experiment) in a log10 scale and methylation density (proportion of cytosines with a probability of methylation greater than 0.5) within transcription factor bound regions. The increasing methylation density has weaker effect on LFY than on the two other TFs. (D-F) Violin plots of DAP/ampDAP signal ratio in a log10 scale as a function of the number of methylated cytosines in the best TF binding site (TFBS) of each bound region. LFY binding is barely affected by the increased number of methylated cytosines. (G-I) Binding site sequence motif for each TF and the methylation effect on each individual position. For LFY, a single half of the symmetric motif is shown. Heatmaps show the Pearson’s correlation coefficient (*r*) between the DAP/ampDAP signal ratio in a log10 scale and the probability of methylation at each position of the best TFBSs. Blank positions have a high false discovery rate (> 5%) and grey indicates positions with less than ten cytosines in the dataset. Correlation are tested on both sides of a symmetric motif (G) or on both strands for non-symmetric motifs (H-I).

### LFY binds to a subset of the closed chromatin regions

Next, we analyzed how *in vivo* factors (including the chromatin state) affect LFY DNA binding. For this, we compared LFY binding *in vitro* and *in vivo* by plotting the coverage of LFY DAP-seq peaks versus that of LFY ChIP-seq peaks. LFY ChIP-seq was obtained from *35S::LFY* seedlings or floral meristems (Sayou et al., 2016; Goslin et al., 2017). This analysis identified genome regions well bound in both experiments (Figure 2A; Supplemental Figure 7A; colored in light purple to red). However, it also highlighted the existence of regions much better bound *in vivo* (ChIP-specific regions colored in deep purple) or *in vitro* (DAP-specific regions colored in orange). The existence of ChIP-specific regions indicated that LFY DNA binding might increase due to interactions with *in vivo* factors. The presence of DAP-specific regions indicated that the *in vivo* context inhibits LFY from binding to some genomic regions despite high affinity LFY binding sites are observed in those regions in DAP-seq.

**Figure 2:**
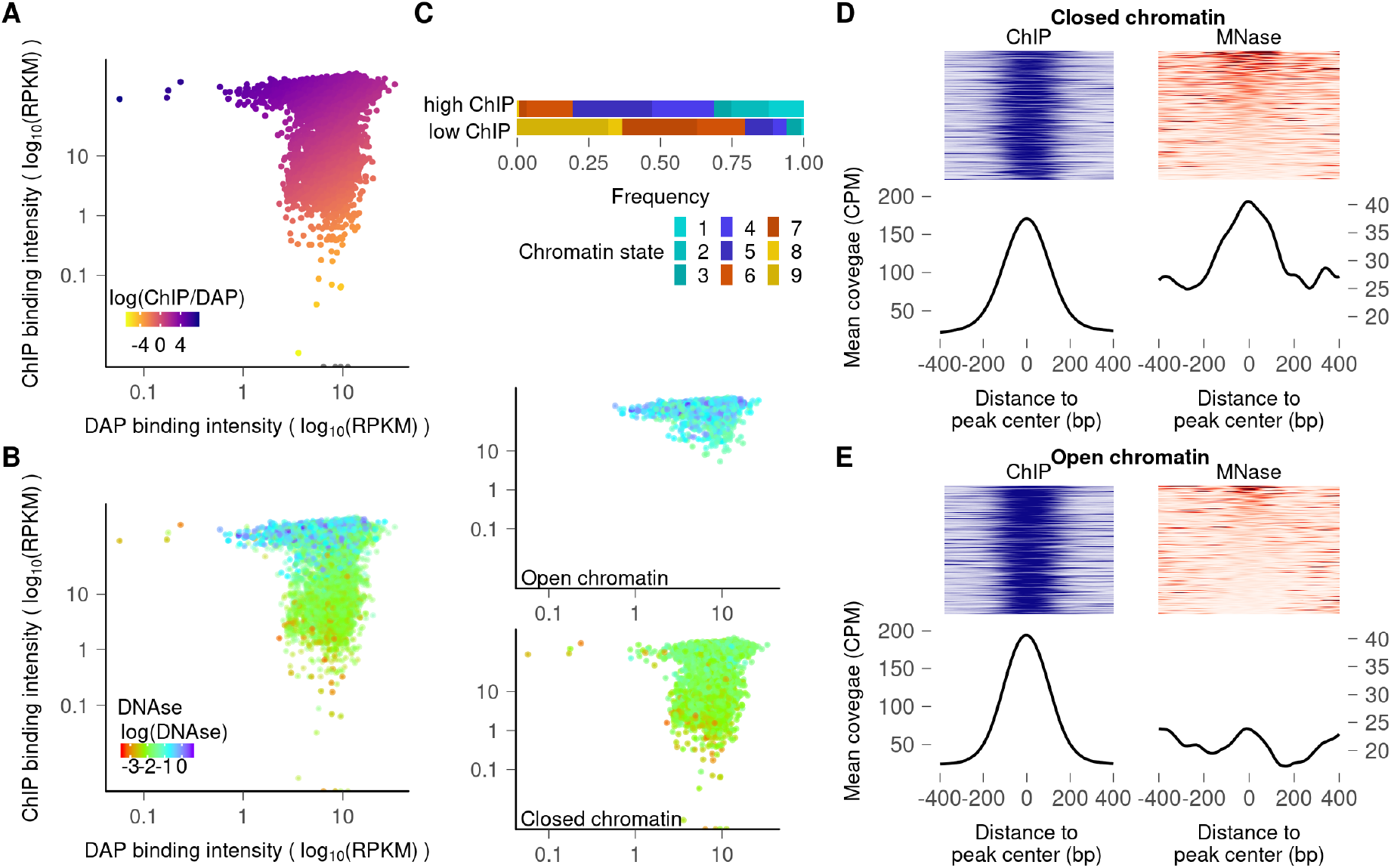
LFY is able to bind nucleosomes in closed chromatin regions. (A) Plots comparing the LFY binding intensities in ChIP-seq vs DAP-seq experiments. Heat map is based on the ChIP-seq/DAP-seq intensity ratio. (B) Overlay of DNaseI signal (heat map) on LFY bound regions, with DAP-seq (X-axis) and ChIP-seq (Y-axis) peak coverages. The two panels on the right show the same regions split into open (upper panel) and closed (lower panel) chromatin states. (C) Distribution of chromatin states 1 to 9 according to (Sequeira-Mendes et al., 2014) for the first and last decile of LFY bound regions based on ChIP-seq signal. (D-E) MNase signal around ChIP-seq peak centers in closed (D) or open (E) chromatin regions. Upper panels show ChIP-seq and MNase-seq coverage for each peak ordered based on MNase-seq signal. Lower panels represent the mean coverage.

To understand whether chromatin conformation could play a role in this inhibition, we analyzed the chromatin state of each region using DNaseI-seq data obtained in two-week-old seedlings (Zhang et al., 2012), a high DNaseI-seq signal being indicative of an open region (Figure 2B; Supplemental Figure 7B). We found that many of the DAP-specific regions have a low DNaseI signal, typical of closed chromatin regions. This suggests that closed chromatin regions inhibit LFY binding. However, as previously observed (Sayou et al., 2016), a number of regions are bound in ChIP-seq despite low DNaseI signal (right panels on Figure 2B and Supplemental Figure 7B). Overall, this analysis suggests that while the closed chromatin context is generally inhibitory for LFY binding, some closed chromatin regions can still be bound. To analyze what type of closed regions are most likely to be bound, we analyzed the upper and lower deciles of regions ranked based on their ChIP-seq signal, the upper decile contains regions well bound in ChIP whereas the lower has regions poorly bound in ChIP (but bound in DAP). The distribution of nine chromatin states (as defined in the literature (Sequeira-Mendes et al., 2014)) changes drastically between the two deciles (Figure 2C; Supplemental Figure 7C). Chromatin states 7, 8, and 9 (that is the most compacted and includes heterochromatin) are unlikely to be bound whereas states 1-5, which represent closed regions but closer to gene units or targets of Polycomb repression (state 5) are more frequently found in LFY bound regions, showing that closed regions are not all equivalently contacted by LFY depending on their functional category.

As closed chromatin regions are often occupied by nucleosomes, and since *in vivo* data suggests that LFY might be able to bind some of these regions, we wondered whether LFY binding was compatible with the presence of nucleosomes. To test this, we compared the position of LFY ChIP-seq peaks with that of nucleosomes (based on MNase-seq data (Zhang et al., 2015)). We found that nucleosomes were indeed enriched at the center of LFY ChIP-seq peaks in closed regions (Figure 2D; Supplemental Figure 7D), but not in open ones (Figure 2E; Supplemental Figure 8E), suggesting that LFY might be able to directly bind nucleosomal DNA *in vivo*. The mapping of LFY TFBS in nucleosome-occupied LFY ChIP-seq peaks show a slight enrichment at the center of the nucleosome, around the dyad position which is a site commonly bound by pioneer TFs (Figure 3A; Supplemental Figure 8) (Zaret, 2020). However, since these genomic data are established on mixtures of tissues, they are not sufficient to firmly establish that LFY is indeed able to bind nucleosomal DNA.

**Figure 3:**
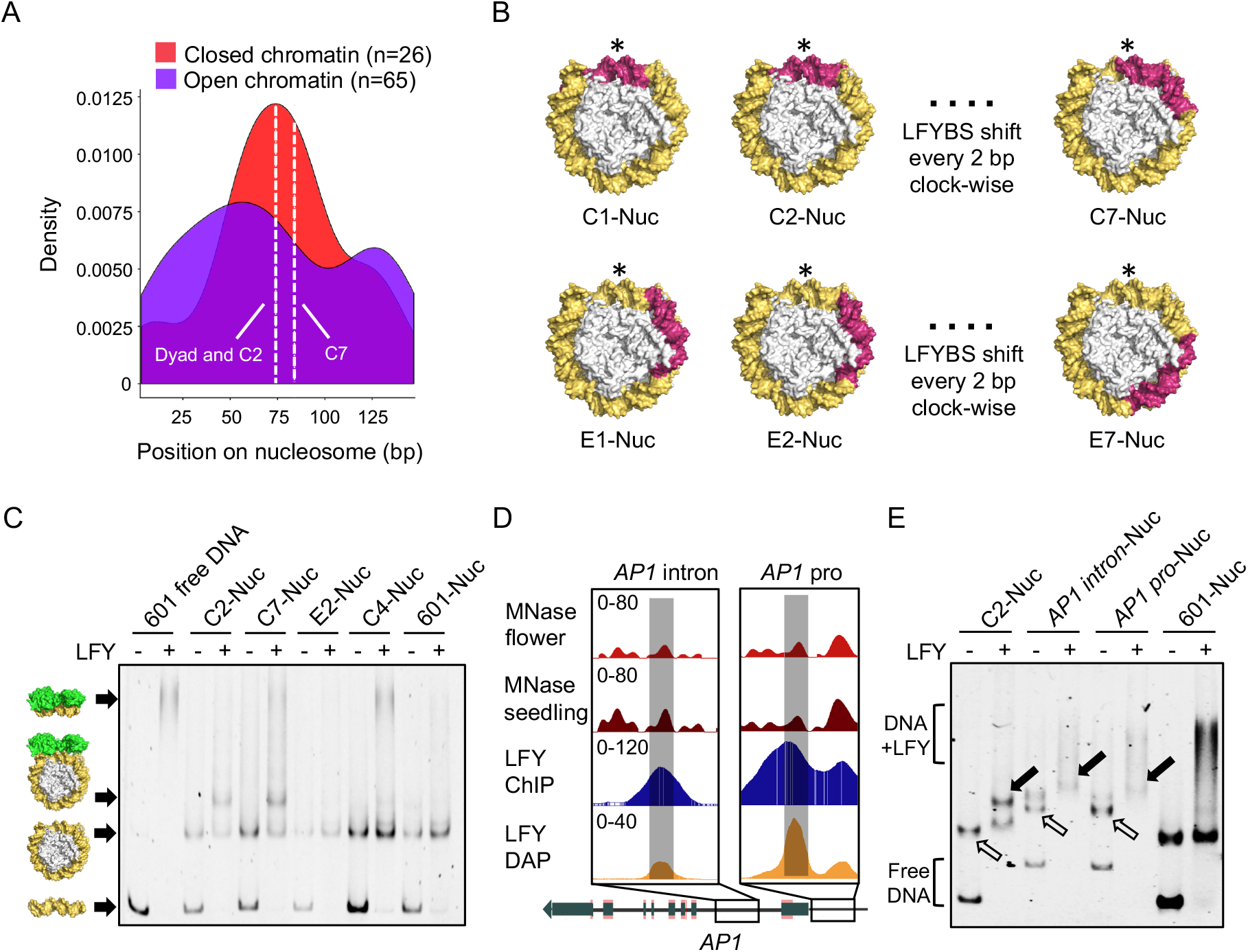
LFY binds nucleosomes *in vitro*. (A) Density plot of the LFY best binding site along a canonical 147-bp nucleosomal sequence in open and closed chromatin contexts for flower tissues. An enrichment for LFY binding sites (LFYBS) around the dyad position (the center of the nucleosomal DNA) is observed in closed chromatin regions. C2 (at dyad) and C7 positions are indicated. Alternative plots for different thresholds for binding sites selection are reported in Supplemental Figure 8. (B) Design of Widom 601 sequences (yellow orange) with a LFYBS (warm pink) inserted at different positions (central C1-C7 (top) and external E1-E7 (bottom)) on nucleosome (PDB: 3UT9 (Chua et al., 2012). * indicates the dyad. (C) Representative EMSA showing LFY binding to 601 nucleosomes with a LFYBS at positions C2 (labelled C2-Nuc) and C7, but not at E2, C4 or 601 nucleosome without a LFYBS (refer to Supplemental Figure 9 for the screening of LFY nucleosomal DNA binding at all other positions). Free DNA (C2 in the first 2 lanes, or present in the nucleosomal preparations) is shifted at the very top of the gels. 601-Nuc is made with wild-type 601 sequence (without a LFYBS): only free DNA is shifted due to non-sequence specific interactions with LFY. Cartoon on the side from bottom to top are free DNA, nucleosome alone, LFY-nucleosome complex and free DNA-LFY complex. (D) Genomic snapshots of LFY DAP-seq, ChIP-seq (seedlings tissue), and MNase-seq (seedlings and closed flower buds) at the *AP1* loci. *AP1* intron and *AP1* pro sequences used to assemble nucleosomes in (E) are highlighted in grey. Both regions are bound in DAP and ChIP, and with well-defined nucleosome signals, lower in floral tissue than in seedlings. (E) EMSA showing LFY binding to nucleosomes assembled with *AP1* intron and *AP1* pro sequences. AP1-intron-Nuc and AP1-pro-Nuc are longer than 601 due to the presence of amplification primers. Note some free 601 DNA is shifted despite the absence of LFYBS in the last lane. The hollow and solid arrows indicate the position of reconstituted nucleosomes and the shifted nucleosomes signals, respectively.

### LFY binds nucleosomal DNA at specific sites *in vitro*

Next, we tested whether LFY has the capacity to bind nucleosomal DNA *in vitro*. We first assembled nucleosomes using the Widom 601 strong nucleosome positioning sequence (Lowary and Widom, 1998; McGinty and Tan, 2015), in which a LFY binding site (LFYBS) was inserted at different positions (C1-C7 around the dyad and E1-E7 farther away) (Figure 3B; Supplemental Table 2). Nucleosomes assembled with a LFYBS at position C2 and C7 were shifted upon addition of LFY, whereas no shift was observed for nucleosomes with a LFYBS at positions C1, C3-C6, E1-E7 or with no LFYBS, demonstrating that LFY binds nucleosomal DNA in a sequence specific manner and only with a LFYBS present at specific positions (C2, located around the dyad, and C7, located one helix turn apart from C2, with the LFYBS exposed to the outer nucleosome surface (Figure 3B and C; Supplemental Figure 9)). Using the same methodology, as a negative control, we tested nucleosomal DNA binding of the TF REGULATOR OF AXILLARY MERISTEMS 1 (RAX1), a direct downstream target of LFY (Chahtane et al., 2013). We found that RAX1 cannot associate with nucleosomes even when its binding site is exposed to the outer nucleosome surface and at the dyad (Supplemental Figure 10), suggesting that RAX1 is unlikely a pioneer TF. We also assembled nucleosomes with two regions of the *AP1* gene, a known early activated LFY target (Parcy et al., 1998; Wagner, 1999; Benlloch et al., 2011). These regions were taken from *AP1* first intron and *AP1* promoter (annotated as *AP1* intron and *AP1* pro, respectively, in Figure 3D). They are both bound by LFY *in vivo* (ChIP-seq (Moyroud et al., 2011; Winter et al., 2011; Sayou et al., 2016; Goslin et al., 2017)) and *in vitro* (DAP-seq in Figure 3D), and with well-defined nucleosome signals from MNase-seq in both seedlings and flower tissues (Zhang et al., 2015) (Figure 3D). We observed that LFY was able to bind to these nucleosomes (Figure 3E and Supplemental Figure 11), showing that LFY nucleosomal DNA binding also occurs within Arabidopsis genomic regions.

### LFY constitutive expression induce changes in nucleosome position

One key characteristic feature of a pioneer TF is to be able to modify the status of closed chromatin regions (Iwafuchi-Doi and Zaret, 2014). To test whether LFY is able to do so, we examined whether it could alter nucleosome positions when ectopically expressed in seedlings. We selected regions that are closed in wild-type seedlings and with a mapped nucleosome (Zhang et al., 2015) and bound by LFY in ChIP-seq and DAP-seq (Figure 4) (Sayou et al., 2016). Among these regions, we focused on the three floral regulators, *AP1* and *AG* (two established LFY targets), and *ULTRAPETALA 1 (ULT1)*, another floral regulator involved in *AG* activation (Moreau et al., 2016). Using Formaldehyde-Assisted Isolation of Regulatory Elements (FAIRE)-qPCR that identifies nucleosome depleted regions, we tested whether ectopic LFY expression (*35S::LFY*) could alter the local chromatin as compared to two-week-old Col-0 seedlings where endogenous LFY is not yet highly induced. We found that LFY ectopic expression induced nucleosome depletion in these regions (Figure 4A). Interestingly, in the *AP1* intron, a region strictly inaccessible in Col-0 seedlings according to DNAseI signal, LFY expression triggers a strong depletion (3-fold, position P1 in Figure 4A). In the *AP1* promoter, a region already largely accessible, LFY induced depletion is more moderate (P2, P3; Figure 4A). As controls, we tested three regions (*Actin2, AT2G38220* and *AT4G22285*) where LFY does not bind *in vivo* and *in vitro* and with poor accessibility in seedlings. We found that their nucleosome status was not altered by LFY ectopic expression (Figure 4B). Taken together, our data suggests that LFY ectopic expression in seedling tissues is sufficient to trigger nucleosome depletion in regulatory regions of some key floral regulators including two LFY target genes.

**Figure 4:**
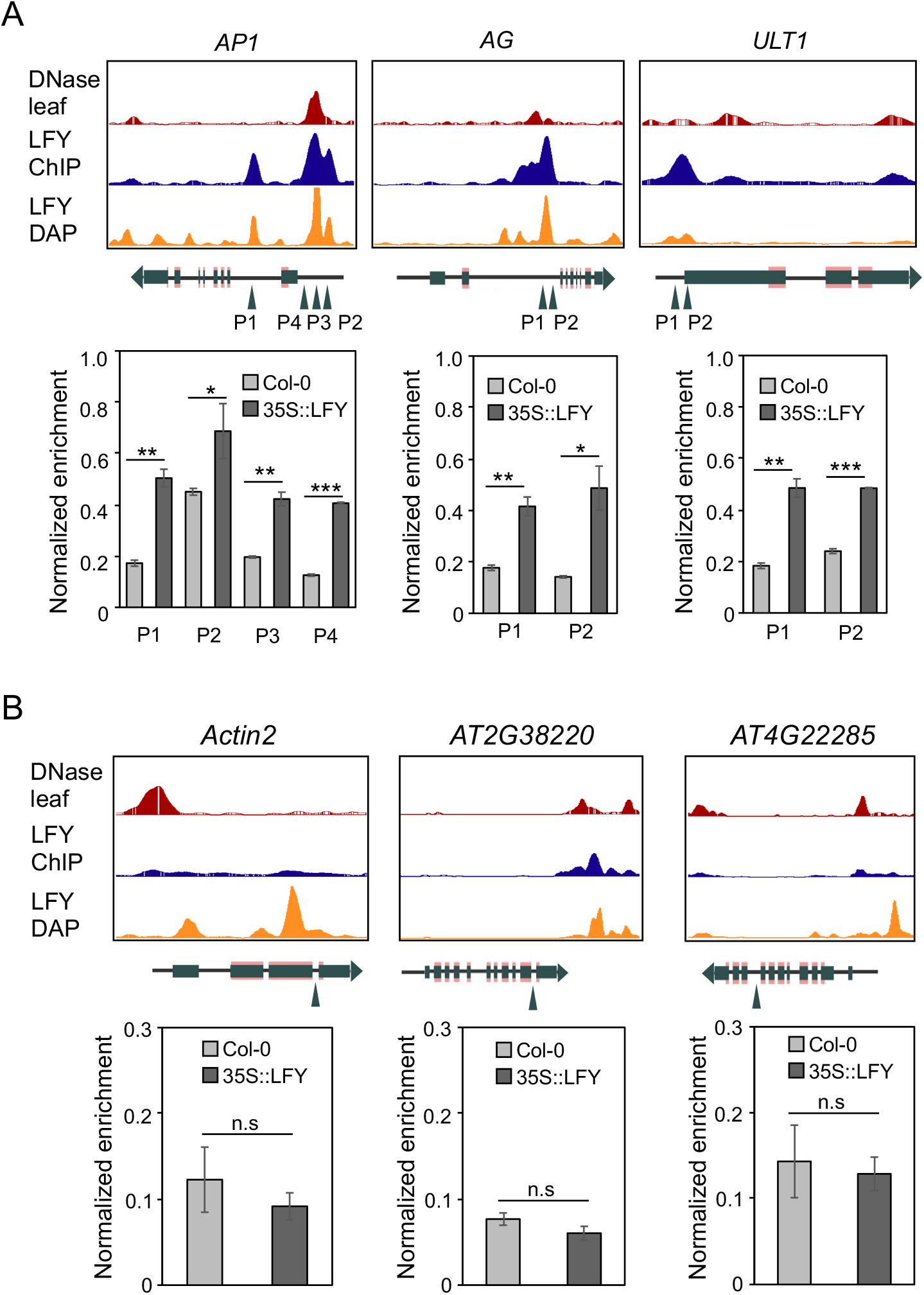
LFY constitutive expression increases chromatin accessibility. (A) (Top) Genomic snapshots of chromatin accessibility (DNaseI-seq from 2-week-old Col-0 seedlings (Zhang et al., 2012)), LFY binding *in vitro* (DAP-seq using genomic DNA from 2-week-old *35::LFY* seedlings), *in vivo* (ChIP-seq of 2-week-old *35::LFY* seedlings (Sayou et al., 2016)) at *AP1*, *AG* and *ULT1* loci. The regions tested in FAIRE-qPCR are indicated by triangle arrows. (Bottom) FAIRE-qPCR of the indicated regions are performed in 2-week-old seedlings of Col-0 (pale gray) and *35S::LFY* (dark gray), respectively. Error bars represent means ± standard deviation. Significance test is performed by one-tailed students’ t-test, *p<0.05, **p<0.01, ***p<0.001, n.s, not significant. (B) (Top) genome browser snapshots of three genomic regions devoid of LFY binding and poorly accessible in 2-week-old seedlings. (Bottom) FAIRE-qPCR on the indicated regions. Significance test is performed as per (A). The FAIRE-qPCR is performed by two biological replicates, with three technical replicates for each. The enrichment is normalized by input DNA in each experiment.

## Discussion

LFY is a master regulator of floral development able to induce the expression of floral organ identity genes that are known to be under repression of Polycomb repressive complex (Goodrich et al., 1997; Turck et al., 2007; Calonje et al., 2008; Kaufmann et al., 2010). In a previous study (Sayou et al., 2016), ChIP-seq data suggested LFY could bind to closed chromatin regions and possibly act as a pioneer factor, although a mechanism was never fully described. By analyzing seedlings constitutively expressing LFY, it was suggested that LFY wild type could bind to regions with a closed chromatin status and that this capacity was strongly impaired upon mutation of the LFY oligomerization sterile alpha motif (SAM) domain (Sayou et al., 2016). Oligomerization would likely increase the DNA-binding affinity of LFY, as has been shown for other TFs able to multimerize (Lai et al., 2020). This increase in DNA binding affinity may be critical for the efficient binding to partially occluded sites in closed chromatin regions, however an increase in binding affinity alone is likely not sufficient to enable recognition of a TFBS in a closed region of chromatin. Insensitivity to methylation state and the presence of nucleosomes are prerequisites to pioneer function, which we investigated here.

Given the high level of DNA methylation found in closed chromatin, it has been hypothesized that some pioneer TFs would be able to bind DNA independently of its methylation status (Zhu et al., 2016; Mayran and Drouin, 2018). Indeed, pioneer TFs from animals, such as Pax7 (Mayran et al., 2018, 7), OCT4 (Yin et al., 2017) and KLF4 (Hu et al., 2013; Liu et al., 2014), are either insensitive or even prefer methylated DNA. The DNA binding of most TFs is inhibited by DNA methylation because the 5-methyl group of methylcytosine often clashes with protein residues that are involved in specific base readout (Yin et al., 2017). Some TFs, however, are not sensitive or even favor methylated DNA because direct hydrophobic interactions form between the methyl group and the TF, as it is the case for homeodomain TFs (Yin et al., 2017) or for some basic leucine zipper TFs (Weber et al., 2019). In this study, we showed that LFY is only mildly sensitive to methylation. We further dissected how methylation impacts LFY binding for each individual position of a canonical LFY binding site (Figure 1).

Consistent with our structural analysis, we showed that the key protein-DNA interactions are not affected by cytosine methylation (Supplemental Figure 6), and even that, at two positions, a methyl group might enhance LFY binding. This computational analysis has the potential to be generalized to all TFs for which DAP and ampDAP data are available. It represents a powerful complement to methylation-sensitive SELEX (systematic evolution of ligands by exponential enrichment) analysis which was used to detect the effect of methylation to TF-DNA binding using randomized DNA sequences (Yin et al., 2017).

Next, we examined whether LFY could bind nucleosomal DNA *in vivo*. Overall, for the majority of regions, a closed chromatin state has inhibitory effect on LFY binding. This is particularly true for heterochromatin regions likely because their high level of compaction totally prevents TF access. However, a subset of closed regions present in the vicinity of genes showed a LFY binding signal in ChIP-seq both in seedlings and floral tissues, consistent with previous observations (Sayou et al., 2016). The limitation of such prior analysis is the heterogeneity of the tissue used: it is conceivable that the observed LFY binding signal would come from the subset of cells where the regions are open. Using *in vitro* reconstituted nucleosomes, we show here that LFY has the capacity to bind nucleosomal DNA. This property is consistent with structural data showing that LFY binds a single side of the DNA and with assays where LFYBS position was either exposed to the outer surface (like C2 or C7) or partially hidden by histones (C1, C3 to C6). Moreover, we found that *LFY* ectopic expression was able to increase nucleosome depletion at several target loci. We examined in particular the *AP1* gene, a known direct target of LFY (Parcy et al., 1998; Wagner, 1999) that is induced immediately after LFY during flower development. LFY binds to two *AP1* regions, its promoter and its first intron. According to DNAseI signal, *AP1* promoter is a region already open in seedlings and with two nucleosomes detected by MNase-seq. This observation likely explains that *AP1* promoter can be induced by LFY already in seedlings and independently of flower formation (Parcy et al., 1998). Here, we observe that LFY ectopic expression is able to induce a mild nucleosome depletion on *AP1* promoter (Figure 4A). The effect on *AP1* intron is more pronounced. This region is closed in seedlings according to DNAseI signal and opens in the flower. Consistently, we observed a strong nucleosome depletion following LFY constitutive expression (Figure 4A). It is thus likely that LFY pioneer property is essential to trigger *AP1* activation through the opening of its closed intronic regulatory region.

Our *in vitro* experiments using reconstituted nucleosomes and LFYBS in various positions further supports LFY’s ability to bind nucleosomal DNA, near the dyad as observed for a subset of animal pioneer TF (Zaret, 2020). Interestingly, LFY binding appears to be fully compatible with the presence of histones, suggesting that LFY may require additional factors for histone displacement. The change in chromatin status following LFY binding might be due to LFY’s capacity to recruit chromatin remodelers such as BRAHMA (BRM) and SPLAYED (SYD) (Wu et al., 2012). These remodelers have been shown to have a very general role (Archacki et al., 2016) and specific mutations altering LFY-BRM or LFY-SYD interactions are needed to fully test this hypothesis.

Our chromatin accessibility assay by FAIRE-qPCR suggests that constitutive LFY expression is sufficient to induce accessibility for a few key floral genes. To fully validate LFY pioneer activity, it would be essential to test its ability to alter chromatin states in the context of floral meristem cells, where the role of LFY is the most prominent and ideally using single cell isolation and next generation sequencing techniques. However, pioneer activity is likely a spectrum of activity in which TFs that play central roles in developmental transitions, such as LFY, are able to fulfill a pioneer role under certain chromatin conditions, in the presence of specific cofactors (Zaret, 2020) and/or for a few distinct loci (Li et al., 2019). Taken together, our *in vitro* and *in vivo* results demonstrate the essential properties of pioneer TFs-the competence to bind closed chromatin and the ability to trigger subsequent opening of these closed regions-are properties of LFY in the context of at least a few key floral regulatory targets. While this manuscript was in preparation, a preprint also describing LFY as a pioneer TF was made available on the bioRxiv server (Jin et al., 2020).

## Materials and methods

### DAP-seq and ampDAP-seq

#### Plasmid construction

Full-length LFY (AT5G61850.1, 420 residues) coding sequence was PCR amplified and cloned into pTnT vector (Promega) with an N-terminal 5XMyC tag to generate construct pTnT-5MyC-LFY.

#### Construction of input libraries for DAP-seq and ampDAP-seq

The input library of ampDAP-seq was PCR amplified from Col-0 genomic DNA (sheared to average size of 200-500 bp by sonication) and constructed according to published protocol (O’Malley et al., 2016; Bartlett et al., 2017; Lai et al., 2020). For input library of DAP-seq, the genomic DNA that retains native methylation pattern was extracted using phenol-chloroform from 2-weeks-old seedlings of *p35S::LFY* line (pCA26 #15) (Sayou et al., 2016) grown on 0.5x Murashige and Skoog medium in long-day conditions.

#### DAP-seq and ampDAP-seq

LFY protein was produced using an *in vitro* transcription/translation system, TnT^®^ SP6 High-Yield Wheat Germ Protein Expression System (Promega L3260), according to the manufacturer’s instructions. In brief, 2 μg input plasmid (pTnT-5MyC-LFY) was used in a 50 μl TnT reaction with 2-hr incubation at 25 °C. DAP-seq was carried out according to published protocol with minor modifications (O’Malley et al., 2016; Bartlett et al., 2017). Briefly, the 50 μl TnT reaction producing LFY was combined with 50 μl IP buffer (20 mM Tris, pH7.5, 150 mM NaCl, 1mM tris(2-carboxyethyl)phosphine, 0.005% NP-40, and proteinase inhibitor (Roche)) and mixed with 20 μL anti-MyC magnetic beads (Merck Millipore). Following 1 hr incubation at 4 °C, the anti-MyC magnetic beads were immobilized and washed three times with 100 μL IP buffer. While the protein was still bound on anti-c-Myc magnetic beads, 50 ng DAP-seq or ampDAP-seq input library pre-ligated with Illumina adaptor sequences was added. The DNA binding reaction was incubated at 4 °C on a rotor for 90 mins, and then washed 6 times using 100 μL IP buffer. The bound DNA was heated to 98°C for 10 min and eluted in 30 μl EB buffer (10 mM Tris-Cl, pH 8.5). The eluted DNA fragments were PCR amplified using Illumina TruSeq primers for 20 cycles, and purified using AMPure XP magnetic beads (Beckman). The libraries were quantified by qPCR using NEBNext Library Quant Kit for Illumina following manufacturer’s instructions. Individual libraries were pooled with equal molarity, and sequenced on Illumina HiSeq (Genewiz) with specification of paired-end sequencing of 150 cycles. Each library obtained around 10 to 20 million raw reads. Both DAP-seq and ampDAP-seq were performed in triplicates.

### Bioinformatic analyses

#### Reads processing and peak calling

DAP-seq and ampDAP-seq read processing and peak calling was performed as previously described (Lai et al., 2020). Briefly, reads were checked and cleaned using FastQC (http://www.bioinformatics.babraham.ac.uk/projects/fastqc/), and NGmerge (Gaspar, 2018) and mapped with bowtie2 (Langmead and Salzberg, 2012) onto the TAIR10 version of the *A. thaliana* genome (www.arabidopsis.org), devoid of the mitochondrial and the chloroplast genomes. The duplicated reads were removed using the samtools rmdup program (Li et al., 2009). The resulting alignment files for each sample were input to MACS2 (Zhang et al., 2008) to call peaks using the input DNA as control. Consensus peaks between replicates (when available) were defined using MSPC (Jalili et al., 2015) (*P*-value cutoff = 10^-4^) for each experiment (DAP-seq, amplified DAP-seq and ChIP-seq from (Goslin et al., 2017)). Each consensus peak was scanned for possible subpeaks, split into several peaks if needed and the peak widths were then normalized to ± 200 bp around the peak maximum. For all the resulting consensus peaks, a normalized coverage was computed as the mean of the normalized read coverage for each replicate (when replicates were available). Because the DAP-seq and ampDAP-seq experiment had different signal-to noise ratio (Supplemental Table 1) we used the total number of reads in consensus peaks for normalization. This normalized coverage (averaged across replica) defines the binding intensity of a TF at a bound region. The ratio of the binding intensity between DAP and ampDAP defines the DAP/ampDAP signal ratio.

#### Measuring the effect of methylation on TF binding

To measure the effect of cytosine methylation on the TF binding affinity, we tested the correlation between the DAP/ampDAP signal ratio and the methylation levels at 1) the whole bound regions, 2) at the TF best binding site (TFBS) in the bound region and 3) at each position of the TFBS. This approach markedly differs from previous analyses (O’Malley) where the change in binding affinity averaged across all binding sites was contrasted at methylated versus non-methylated regions in DAP and ampDAP experiments separately. By using the DAP/ampDAP signal ratio as a function of methylation levels at a same locus, our approach account for the variability across binding sites and better controls for differences in signal-to-noise ratio between ampDAP and DAP experiments. TFBSs were searched in bound regions using a position weight matrix (PWM) constructed for each TFs using MEME (Machanick and Bailey, 2011). The probability of cytosines methylation was taken from (Zhang et al., 2016). Methylation density (the number of methylated cytosines in a bound region) was defined as the number of cytosines with a probability of methylation greater than 50%. Association between the relative binding intensity and methylation levels was assessed using Pearson’s correlation tests from R package. The effect of methylation on LFY binding was compared to that of two others TFs (AT1G19210 and AT3G10030) for which DAP-seq and ampDAP-seq samples were available from (O’Malley et al., 2016).

#### ChIP-seq versus DAP-seq binding affinity comparison

The ChIP-seq datasets used were obtained from (Sayou et al., 2016) (1,954 peaks, two-week-old seedlings *35S:LFY* tissue), or re-computed (see above) from (Goslin et al., 2017) (884 peaks, inflorescence tissue of *35S:LFY-GR ap1 cal*). Only the first 3,000 DAP-seq bound regions with lowest p-value were considered. Regions bound either in DAP-seq or in ChIP-seq were merged in a single bed file. When peaks overlapped for more than 80% of their respective length, they were considered as “common” and resized to create a common peak. Coverages of pooled bound regions were retrieved for both datasets.

#### DNA accessibility

Closed and open chromatin regions were defined according to leaf DNaseI-seq dataset from (Zhang et al., 2012). DNaseI-seq coverage was computed on the DAP and ChIP pooled peaks and a peak was classified as open or closed following DHs regions from Zhang et al.

#### Chromatin state

9 chromatin states was taken from (Sequeira-Mendes et al., 2014). Those chromatin states were then crossed with our pooled peaks from ChIP-seq and DAP-seq and separated in deciles along the ChIP-seq to DAP-seq coverage fold ratio (CFR).

#### Nucleosomes

MNase-seq defined genomic positions of nucleosomes in leaf were retrieved from (Zhang et al., 2015). Custom python scripts were used to compute MNase-seq coverage for DNase-defined closed and open bound regions. Peaks were then extended to 800 bp, around the maximum, and sorted on their center (+/-100 bp around the center).

#### Position of LFYBS on nucleosomes

LFY ChIP-seq bound regions were considered to be in an open chromatin state if they overlap by more than 50% with a DNase peak in flower tissues, else they were considered to be in closed chromatin. MNase data from flower tissues was crossed (using bedtools intersect -f 0.8 (Quinlan and Hall, 2010)) with ChIP-seq peaks to retain nucleosomes that are 80% within a LFY bound region. The resulting nucleosome sequences, plus half of a LFYBS (i.e. 9bp) at both sides, were screened for LFYBS using a custom LFY position weight matrix (Sayou et al., 2016). We then counted the number of LFYBS, taking the center of the motif as reference, present along the 147 bp sequence of canonical nucleosome.

### Protein structural analysis

The structure coordinates of LFY (accessions of 2vy1 and 2vy2 (Hamès et al., 2008)) are taken from protein data bank (https://www.rcsb.org). The cytosine methylation mutation was done using “Builder” option from the PyMOL GUI (www.pymol.org), all visualization was prepared using PyMOL.

### Protein purification

The protein AtLFYΔ40 was produced in *E. coli* Rosetta2 (DE3) strain (Novagen). Cells were grown in Luria-Bertani medium supplemented with Kanamycin (50 μg/ml) and Chloramphenicol (34 μg/ml) at 37 °C under agitation up to an optical density of 600 nm of 0.6. Betaine (2 mM) was added and cultures were shifted to 18 °C for 1 h before addition of 0.4 mM isopropyl β-D-1-thiogalactopyranoside. After overnight growth at 18 °C, cells were pelleted by centrifugation. Pellets corresponding to 0.5 l culture containing the recombinant protein were sonicated in 50 ml of lysis buffer (20 mM Tris–HCl pH 8.5, 1 mM TCEP) supplemented by one protease inhibitor cocktail tablet Complete EDTA-free (Roche) and centrifuged for 30 min at 20,000 g 4°C. The clear supernatant was transferred on a column containing 1 ml Ni-Sepharose High Performance resin (GE Healthcare), washed two times with lysis buffer containing 20 mM and 40 mM imidazole, respectively, and eluted with lysis buffer containing 300 mM imidazole. Eluted fractions were immediately diluted three times in buffer without imidazole and dialysed overnight. Protein concentrations were determined by SDS-PAGE, using a BSA standard curve run on the same gel.

Recombinant histones were produced according to published protocols (Shim et al., 2012). The coproduction of the *Xenopus laevis* four core histones was done using a pET29a polycistronic plasmid containing the four core histones. The histoneH2A was tagged with N-terminal hexahistidine (his6)-tag with a thrombin cleavage site. Histone H4 was tagged with a C-terminal His6-tag preceded by a thrombin cleavage site. This plasmid was transformed in BL21(DE3)pLysS bacteria. Cells were grown in Luria-Bertani medium supplemented with Kanamycin (50 μg/ml) and Chloramphenicol (34 μg/ml) at 37 °C under agitation up to an optical density of 600 nm of 0.6. were shifted to 18 °C for 1 h before addition of 0.4 mM isopropyl β-D-1-thiogalactopyranoside. cells were pelleted by centrifugation. Pellets containing the recombinant protein were sonicated in 50 ml of lysis buffer (20 mM Tris–HCl pH 7.5, 2 M NaCl 1 mM TCEP) supplemented by one protease inhibitor cocktail tablet Complete EDTA-free (Roche) and centrifuged for 30 min at 18,000 g 4°C. The clear supernatant was transferred on a column containing 3 ml Ni-Sepharose High Performance resin (GE Healthcare), washed two times with lysis buffer containing 30 mM and 50 mM imidazole, respectively. Elution was performed with lysis buffer containing 300 mM imidazole. Fraction were analyzed on 18% SDS-PAGE. Thrombin digestion was carried out by adding purified thrombin in 25:1 mass ratio and incubating the samples at room temperature for 4 hours. The digestion was confirmed by SDS-PAGE. Digested histones were then concentrated by centrifugation using amicon membrane (MW50KDa) at 4°C et 4,000xg during 20 min intervals, with gentle mixing between each centrifugation. The concentrated sample was then injected onto a Superdex 200 10/300 column in lysis buffer. The histone octamer peak was eluted at an elution volume of 12.8 ml. the peaks fractions were pooled and concentrated at a concentration of 1.84 mg/ml and aliquoted flash-frozen in the presence of 50% glycerol.

### Nucleosome reconstruction

#### DNA sequences used for nucleosome reconstruction

To test the interaction between AtLFYΔ40 and nucleosomes we reconstituted nucleosomes with the 601 sequences and different 601 sequences containing a LFY binding site at different positions (C1-C7 and E1-E7, Figure 3B). As a control, we also tested RAX1 nucleosomal DNA binding using 601 sequence containing a RAX1 binding site (C1-C7, Figure 3B). All cloned in a pUC57 plasmid (sequences see Supplemental Table 2 and 3). Sequences of an *AP1* intronic region and *AP1* promoter region (Figure 3D) were also cloned in pUC57 and used for nucleosomal reconstitution (sequences see Supplemental Table 2).

All fragments used for nucleosome reconstruction were amplified by PCR using Invitrogen primer labelled with CY5 fluorophore. The resulting fragment was then checked by 1% agarose gel and purified with the Monarch^®^ DNA Gel Extraction Kit (New England Biolabs). The purified fragment was precipitated by ethanol precipitation and resuspended in buffer (25 mM Tris, pH 7.5, 2 M NaCl and 1mM TCEP) and adjusted at a concentration of 200 ng/μl.

#### Nucleosome reconstruction

The nucleosomes assembly was performed by salt dilution method (Okuwaki et al., 2005). Briefly, DNA of interest and recombinant histone octamer were mixed at a molar ratio (DNA / histones) between 1/1 and 1/1.2 in a solution of 25 mM Tris pH 7.5 2M NaCl 1mM TCEP. This mix was incubated at 30°C for 20 min. The reaction was serially diluted to 1.5, 1, 0.8, 0.6, 0.5, 0.4, 0.3, and 0.2 M NaCl using buffer 25 mM Tris pH7.5 1mM TCEP with 20 min incubation at 30°C for each dilution step. The reconstitution was confirmed by native gel analysis.

#### Electrophoretic Mobility Shift Assay

Nucleosomes of interest were incubated with 500 μM AtLFYΔ40 in buffer (25 mm Tris pH7.5 200mM NaCl 1mM TCEP 10% glycerol 0.1 mg/ml BSA 0,12 mg/ml herring and salmon sperm DNA for 1 hour at room temperature. The different complex was separated on 5% non-denaturing polyacrylamide gels run in 0.5X Tris-borate –EDTA (TBE) buffer. Gels run for one hour at 4°C at 120 V. Complexes were visualized with Cy5 – exposition filter (ChemiDoc MP Imager; BIO-RAD).

### Formaldehyde-Assisted Isolation of Regulatory Elements (FAIRE)-qPCR

#### Site selection for FAIRE-qPCR

To test the effect of LFY binding on nucleosomes, LFY ChIP-seq peaks (Sayou et al., 2016) with a nucleosome in leaf tissue (overlap >= 50%) and no nucleosomes in floral tissue were selected (Zhang et al., 2015). Selected peaks were then attributed to nearby genes (peaks within 3 kb upstream and 1 kb downstream; overlap >= 80%). In those regions, we selected two known LFY targets, *AP1* (AT1G69120) and *AG* (AT4G18960), and *ULT1* (AT4G28190), another floral regulator. We applied a similar method to select three control regions that are not bound by LFY and are occupied with nucleosomes in leaf and floral tissue (AT3G18780 (Actin2), AT2G38220 and AT4G22285).

#### FAIRE-qPCR

FAIRE assays were performed on two-week-old seedlings of Arabidopsis thaliana ecotype Columbia (Col-0) background and 35S::LFY. Seeds were surface-sterilized by treatment with bayrochlore, washed, then sown in sterile half-strength MS medium and placed for 2–4 days at 4 °C to obtain homogeneous germination. Plants were grown in square petri dishes in growth chambers at 20 °C under long-day (16 h of light) conditions. 1g of plant material was then crosslinked in 1% (v/v) formaldehyde at room temperature for 15 min using vacuum infiltration. Crosslinking was quenched by adding glycine solution to a final concentration of 0.125M under vacuum infiltration for 5 minutes. The crosslinked plantlets were ground into powder using liquid nitrogen and nuclei were isolated using Nuclei Isolation Buffer (0,25M Sucrose, 10mM Tris-HCl pH8, 10mM MgCl2, 1% Triton X-100, 5mM β-mercaptoethanol, and proteases inhibitors) and then resuspended in 1ml of FAIRE Lysis Buffer (0,1% SDS, 50 mm Tris-HCl pH 8, 10 mm ethylene diamine tetraacetic acid (EDTA) pH 8). The crosslinked DNA was sheared to an average size of 200 - 300 bp using a Covaris S220 (Peak Power: 175W, cycles/burst: 200, Duty Factory: 20, 4min). Samples were centrifuged for 15 min and 13,000 x g at 4 °C and the supernatant was transferred into new tubes. A 100μl aliquot was used as control DNA and directly treated with 1μl of RNAse A+T1 cocktail enzyme mix (Thermo Fisher Scientific) for 1h at 37°C followed by proteinase K treatment for 4h at 37°C and 6h at 65°C to reverse the crosslinks. The non-de-crosslinked samples were RNAse A+T1 treated as for control DNA and a phenol:chloroform:isoamyl alcohol (25:24:1) extraction was performed to purify DNA and control DNA in a final volume of 100 μL TE buffer (10 mM Tris pH 8, 1mM EDTA). The non-de-crosslinked free DNA samples were then incubated overnight at 65 °C to remove inter-DNA crosslinks. DNA was quantified using Qubit ds DNA HS kit (Thermo Fisher Scientific) and the ratio between nucleosome-free DNA versus total DNA was determined by qPCR analysis using 20ng of template DNA for each reaction.

### Accession Numbers

LFY DAP-seq sequencing data from this article can be found in the NCBI GEO data libraries under accession numbers GSE160013 (token afwnayckrrajfev for reviewers).

## Acknowledgements

We thank K. Kaufmann for discussions. This project was supported by the ANR-DFG project Flopinet (ANR-16-CE92-0023-01) to CZ and FP, the GRAL LabEX (ANR-10-LABX-49-01) with the frame of the CBH-EUR-GS (ANR-17-EURE-0003) to CZ, FP and AS.

## Authors contributions

FP, CZ, RD designed and supervised the project, RBM, JL, AS performed bioinformatics analyses, LG, GV, JLM, HD, ET and EBH performed biochemical analyses, XL performed DAP-seq, YW performed FAIRE supervised by MB and DL, FP, CZ and XL wrote the paper with the help of all authors.

## Supplementary information

**Supplementary Figure 1:**
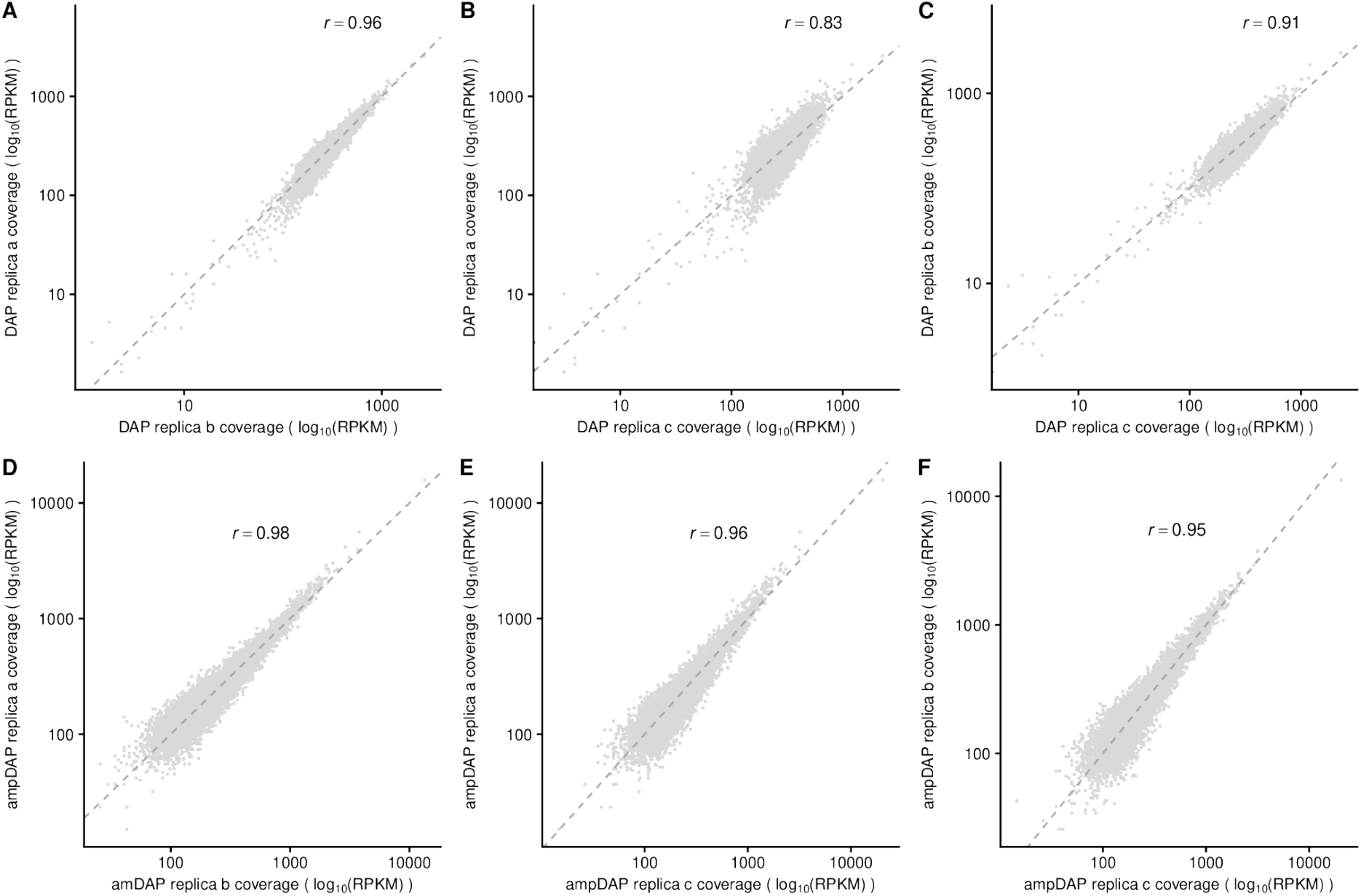
Replicate reproducibility of LFY DAP-seq experiments. Scatter plots of coverage, normalized by the total number of reads in peaks, between three replicates (a,b,c) of DAP-seq (first row) and amplified DAP-seq (second row) experiments, with Pearson’s correlation at top.

**Supplementary Figure 2:**
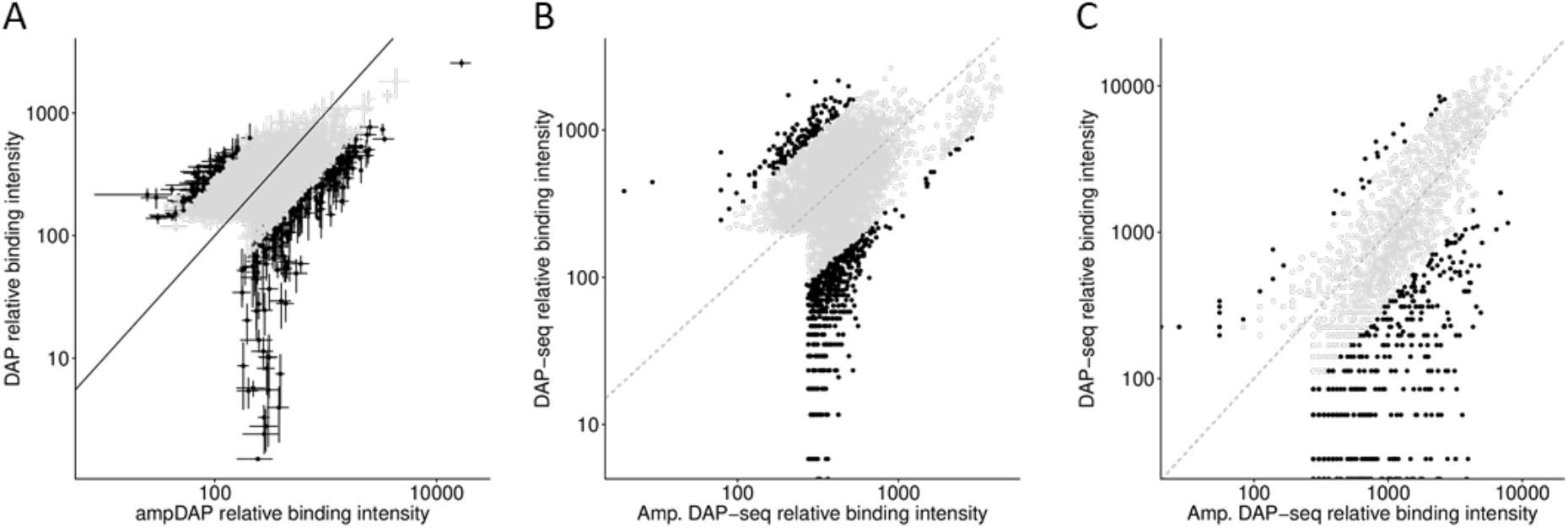
Relative binding intensity for ampDAP-seq and DAP-seq experiments for LFY (A), ERF07(AT1G19210) (B), and a trihelix-domain containing protein (AT3G10030) (C). The relative binding intensity is expressed as the number of reads in the bound region normalized by its length and by the total number of reads in all bound regions. Black and grey separate bound regions with a difference in relative binding intensity of ampDAP versus DAP-seq experiment greater and less than 3-fold. Variability across replicates (available for LFY only) is represented by error bars.

**Supplementary Figure 3:**
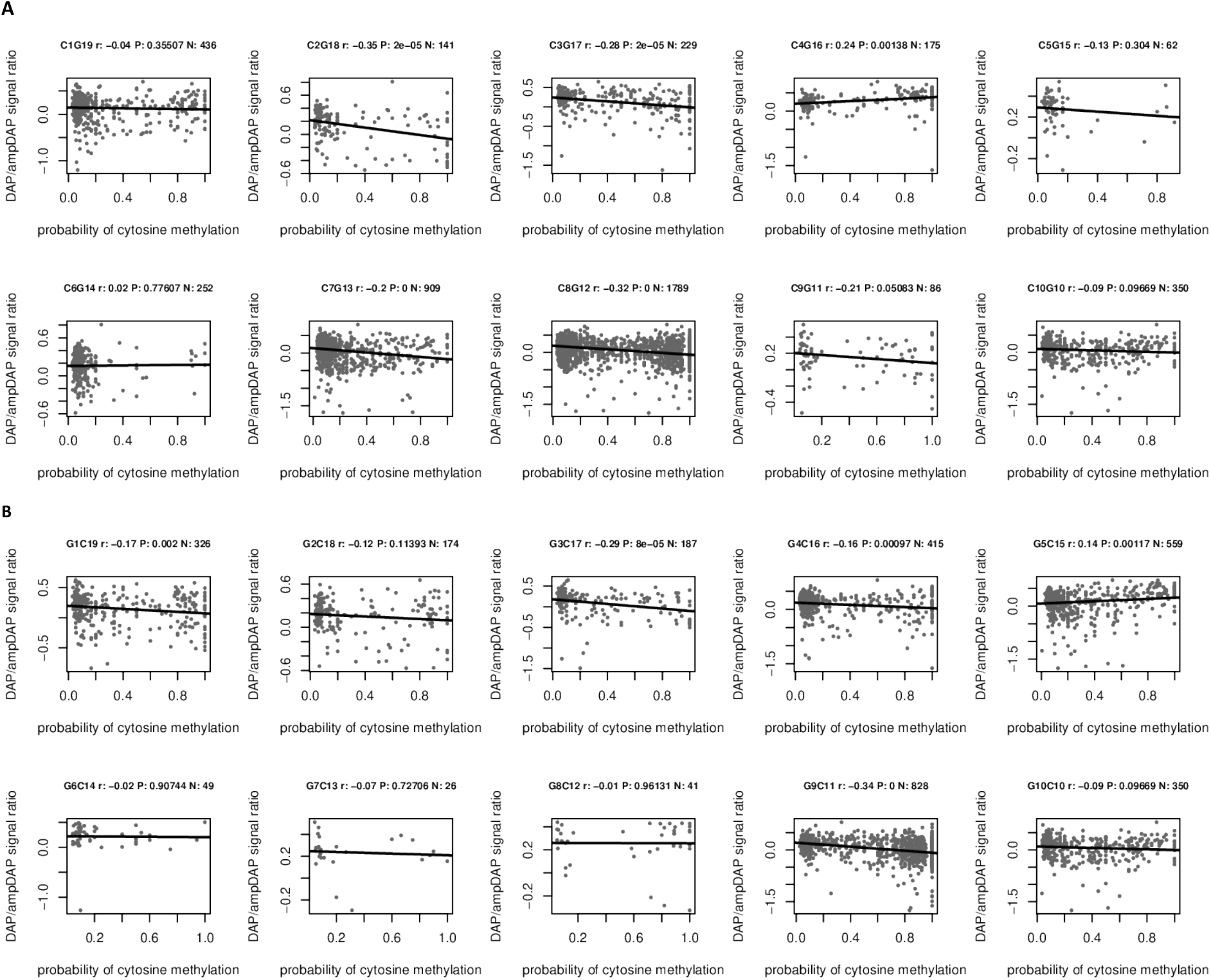
Effect of methylation of each individual position on LFY binding. Relation between methylation probability at a single site in the predicted best LFY binding site and the log10-scaled relative binding site intensity of a DAP-seq versus an ampDAP-seq experiment for LFY on the forward (A) and reverse (B) strand.

**Supplementary Figure 4:**
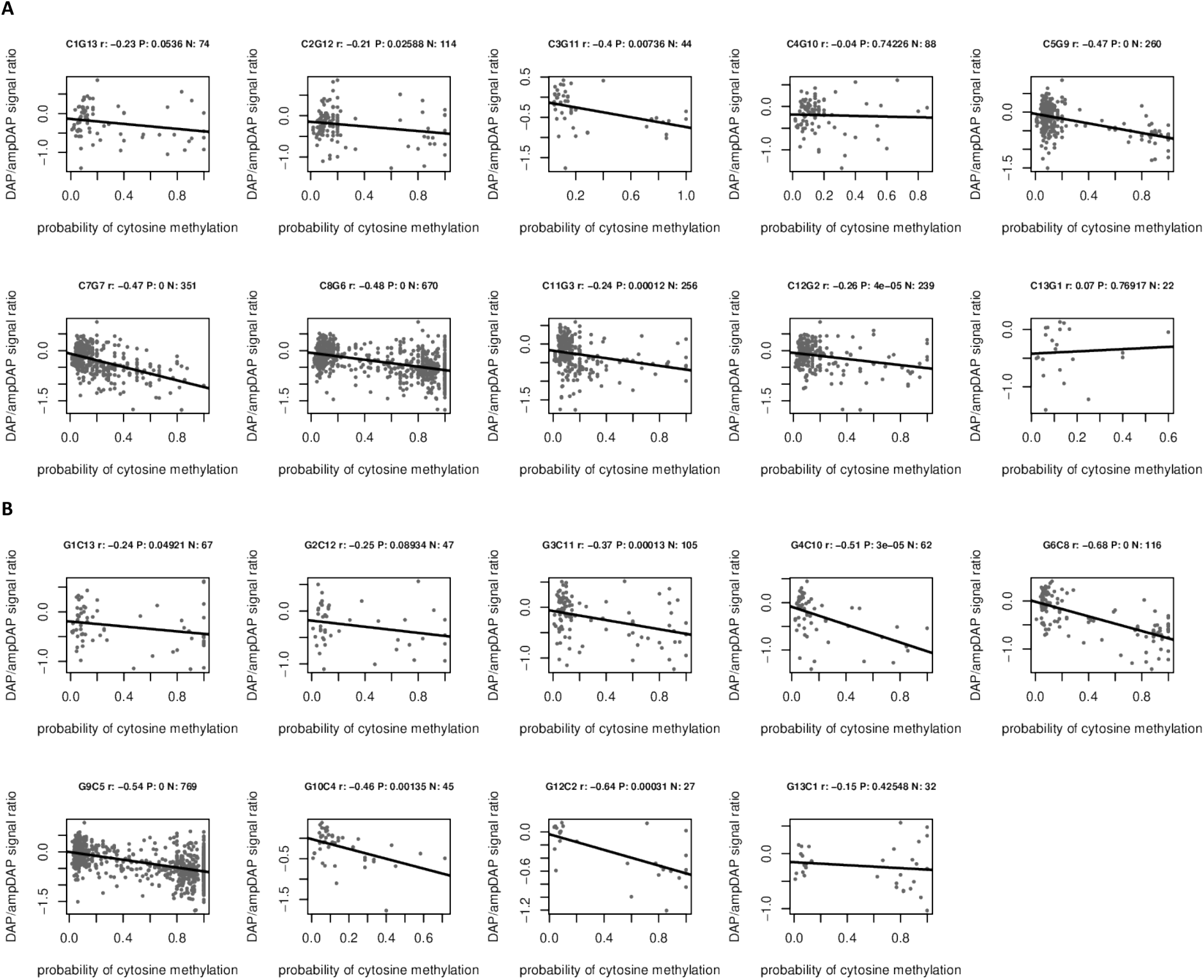
Effect of methylation of each individual position on ERF017 binding. Relation between methylation probability at a single site in the predicted best binding site and the log10-scaled relative binding site intensity of a DAP-seq versus an ampDAP-seq experiment for ERF017 (At1g19210) on the forward (A) and reverse (B) strand.

**Supplementary Figure 5:**
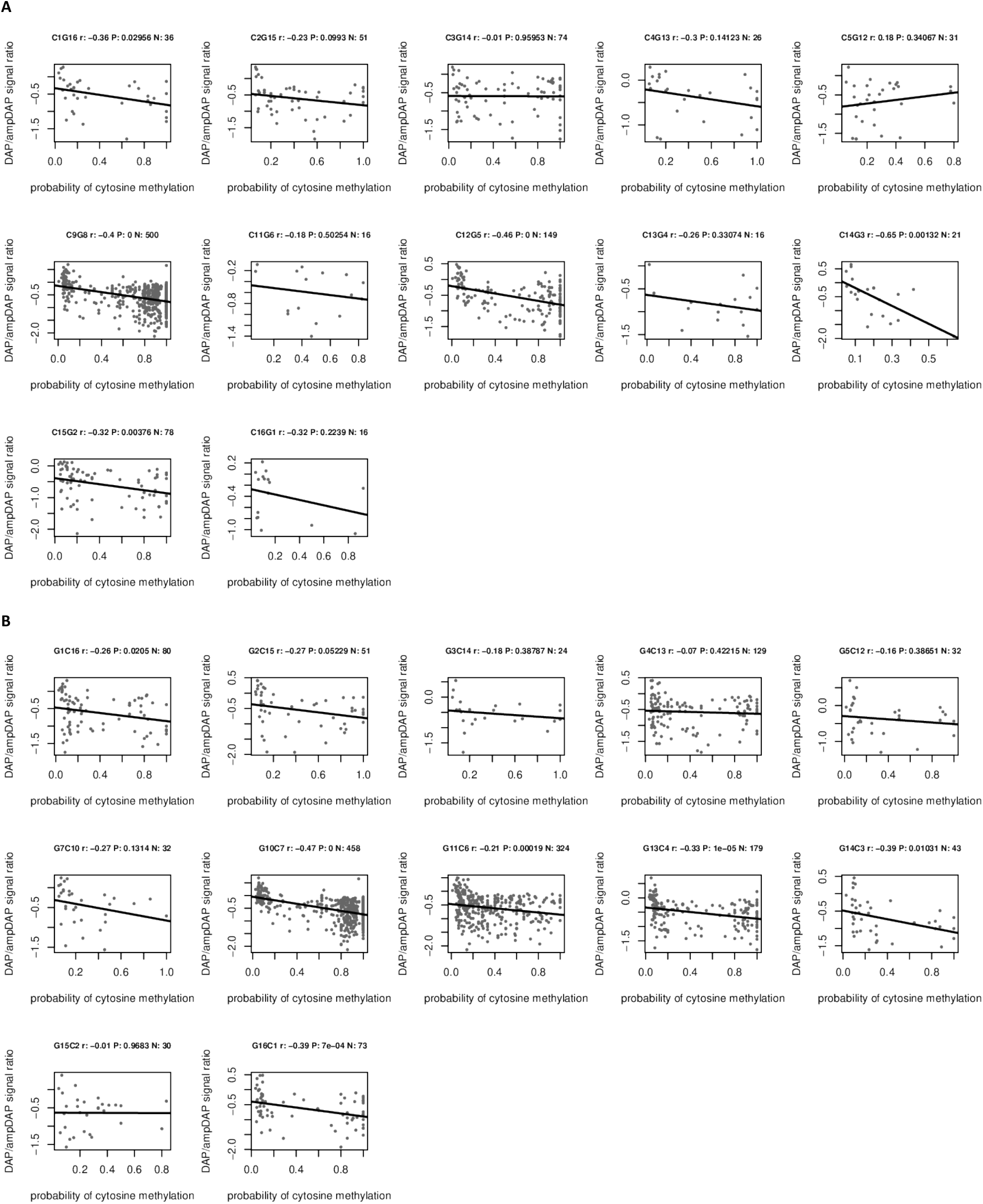
Effect of methylation of each individual position on trihelix-domain containing protein (AT3G10030) binding. Relation between methylation probability at a single site in the predicted best binding site and the log10-scaled relative binding site intensity of a DAP-seq versus an ampDAP-seq experiment for trihelix-domain containing protein (AT3G10030) on the forward (A) and reverse (B) strand.

**Supplementary Figure 6:**
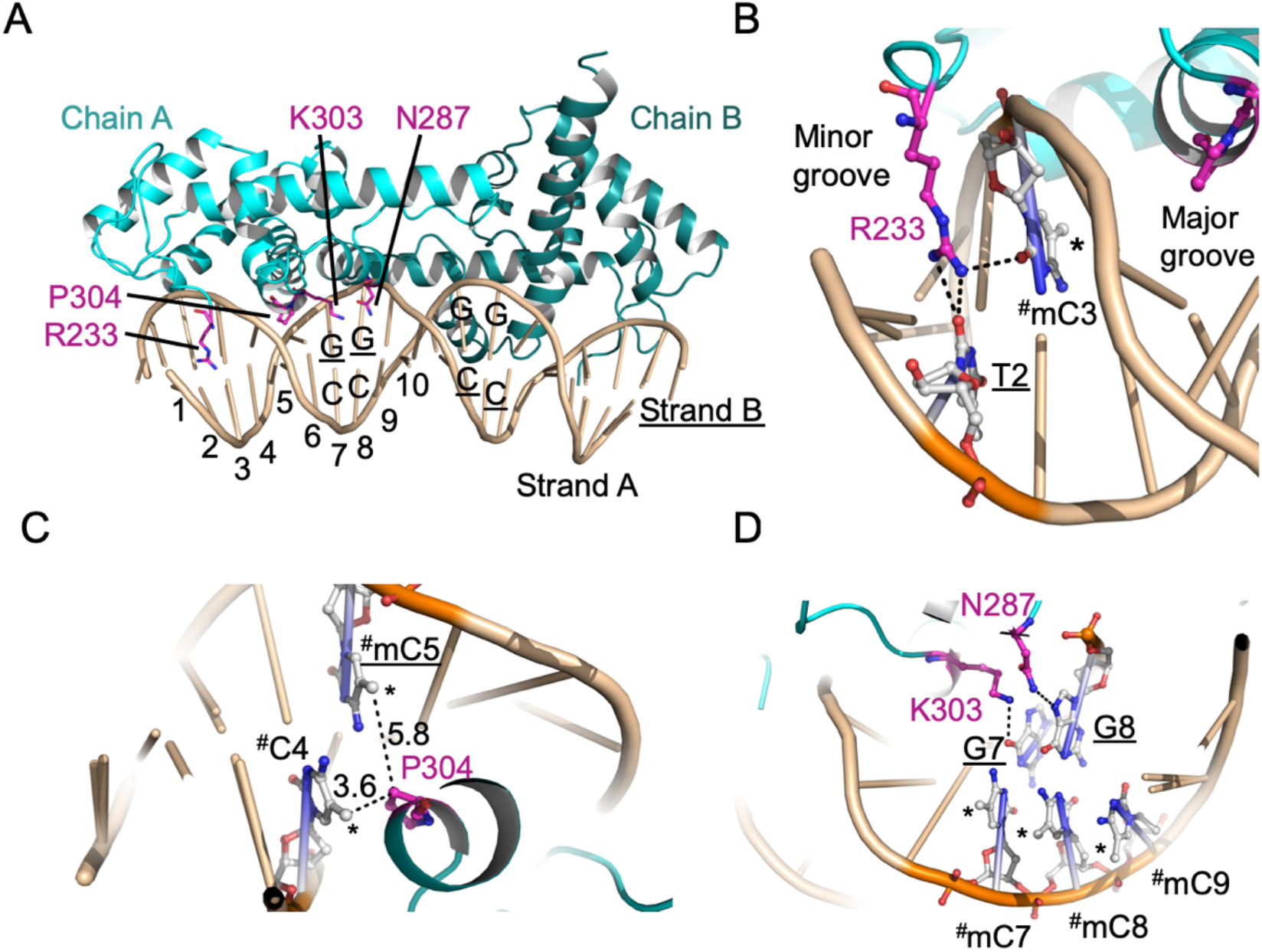
Structural features suggesting that LFY is intrinsically only mildly sensitive to cytosine methylation. (A) Overview of LFY-DBD dimer (Chain A and B) bound to DNA sequence from *AP1* promoter (PDB 2vy1); Residues involved in base readout are highlighted in pink. LFY binding sequence is pseudo-palindromic and composed of two half-sites. One half-site is numbered from 1 to 10 with position 10 being the center of the pseudo-palindrome. The two DNA strands are designated as strand A and strand B (underlined), respectively. The highly conserved CG base pairs in both strands are annotated (bases from strand B are underlined). (B) Base readout residue R233 interacts with bases from position 2 and 3 in the minor groove. C3 is mutated to methylated cytosine, designated as #mC3 (# stands for mutation). Its methyl group facing the major groove, thus would not interfere with R233 base readout interactions. (C) Methylation of cytosine in position 4 (strand A) or position 5 (strand B) helps LFY binding by forming hydrophobic interaction between methyl groups and Cβ of P308; The methyl groups are in close proximity to P304. Distances are measured based on mutation done in the model of PDB 2vy1. (D) C7, C8, and C9 are mutated to mCs. Their methyl groups are distant from N287 and K303 (>8 Å), thus are unlikely to interfere with the base readout interactions. The base readout interactions are indicated by black dash lines. Methyl groups are indicated by *. A base marked with # stands for a mutated base relative to the native structure. It has to be noted that the methylation pattern in the figure is to visualize their relative position to key base readout residues, they might not occur *in vivo*, e.g., the three continuum mC in (D).

**Supplementary Figure 7:**
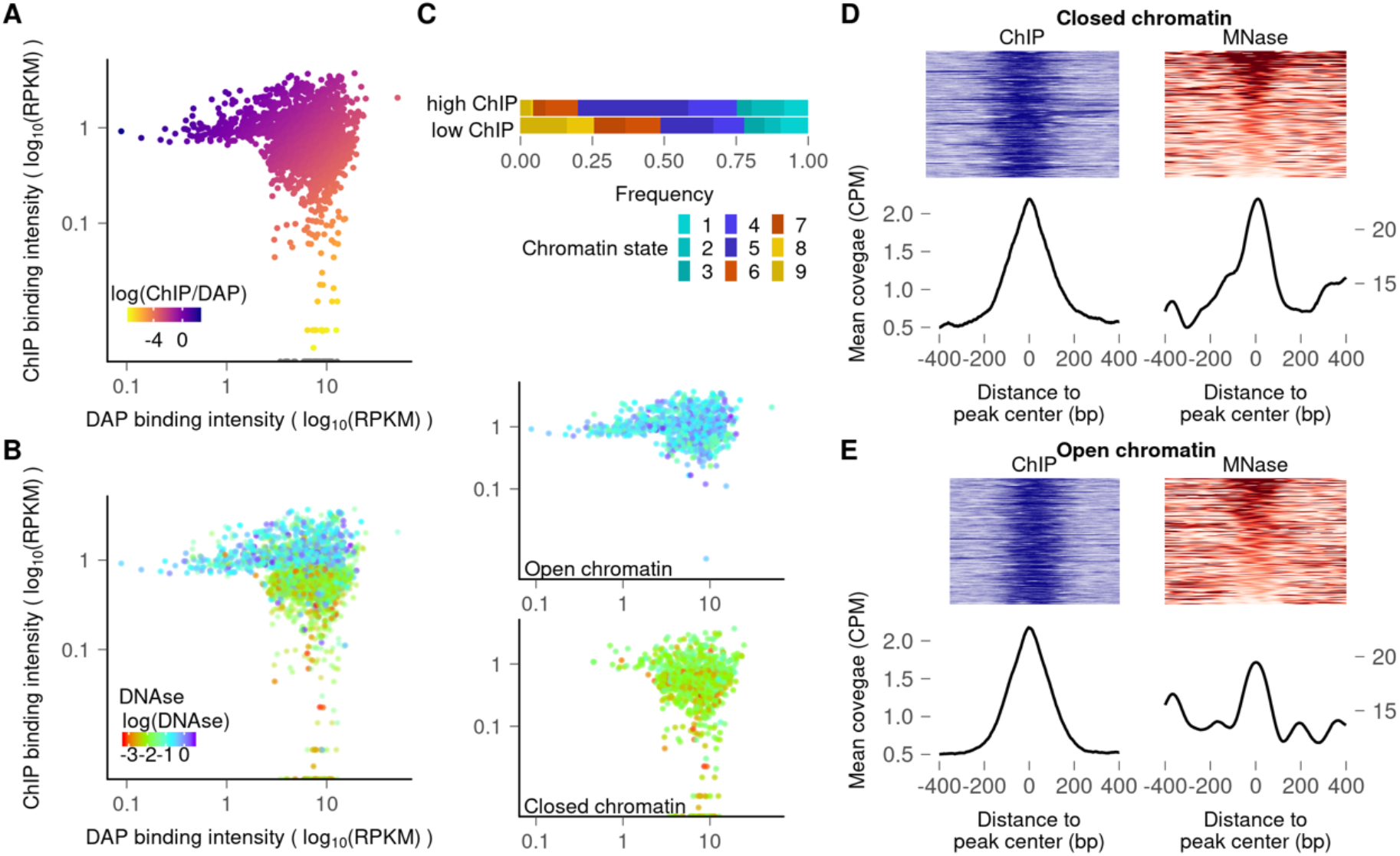
Interaction between LFY and closed chromatin regions in inflorescence tissues. (A) Plots comparing the LFY binding intensities in ChIP-seq (inflorescence tissue of *35S::LFY-GR ap1 cal*; Goslin et al., 2017) vs DAP-seq experiments. The heatmap is based on the ChIP-seq/DAP-seq intensity ratio. (B) Overlay of DNaseI signal (heat map) on LFY bound regions, with DAP-seq (X-axis) and ChIP-seq (Y-axis) peak coverages. The two panels on the right show the same regions split into open (upper panel) and closed (lower panel) chromatin states. (C) Distribution of chromatin states 1 to 9 according to (Sequeira-Mendes et al., 2014) for the first and last decile of LFY bound regions based on the ChIP-seq signal. (D-E) MNase signal around ChIP-seq peak centers in closed (D) or open (E) chromatin regions. Upper panels show ChIP-seq and MNase-seq coverage for each peak ordered based on the MNase-seq signal. Lower panels represent the mean coverage.

**Supplementary Figure 8:**
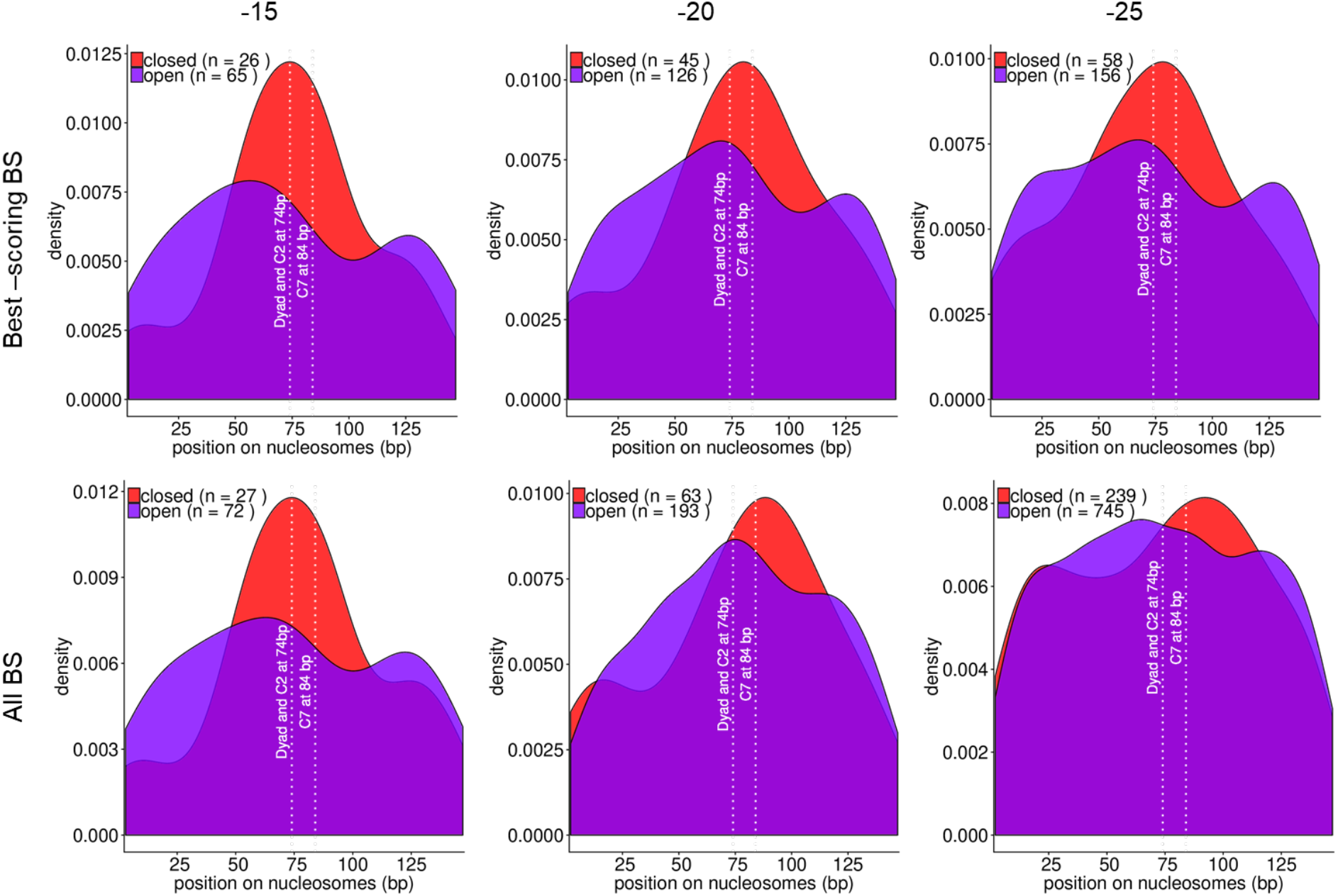
Density plots of LFY binding sites (BS) central position on nucleosomes in open and closed chromatin for flower tissues. Plots are shown for BS with a score greater than −15 (left), −20 (middle), and −25 (right), for best-scoring BS only (top), and all BS above the score threshold (bottom).

**Supplementary Figure 9:**
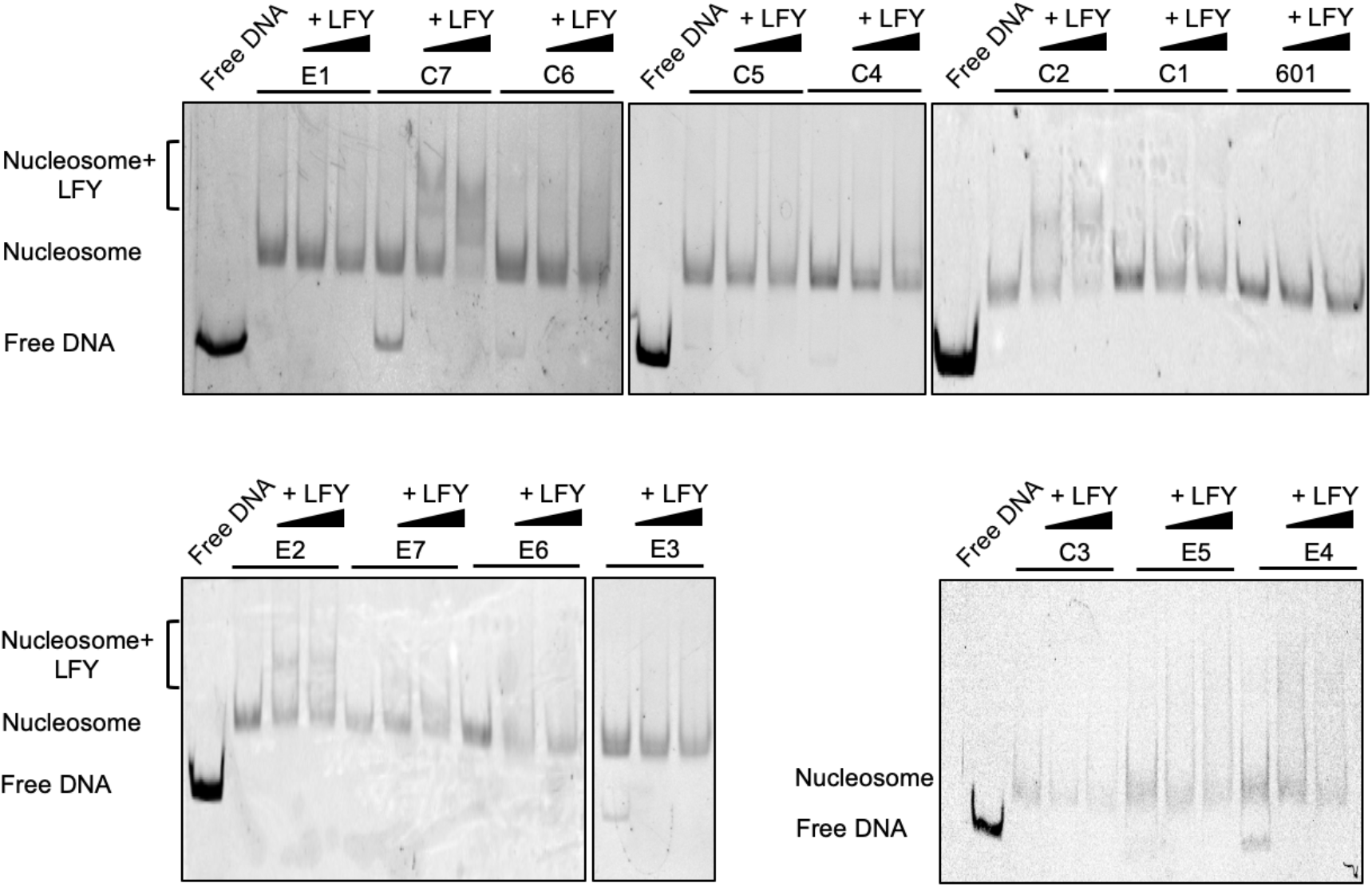
LFY nucleosomal DNA binding assayed by EMSA using 601 sequences with a LFYBS inserted at different positions. The positions of LFYBS in the 601 sequence are illustrated in Figure 3B and their sequences are given in Table S2. Two concentrations of LFY (250 nM and 1 μM) were used. LFY binds to nucleosomes with a LFYBS at C2 and C7 positions; There is also weak binding at E2 but it is weak and less reproducible.

**Supplementary Figure 10:**
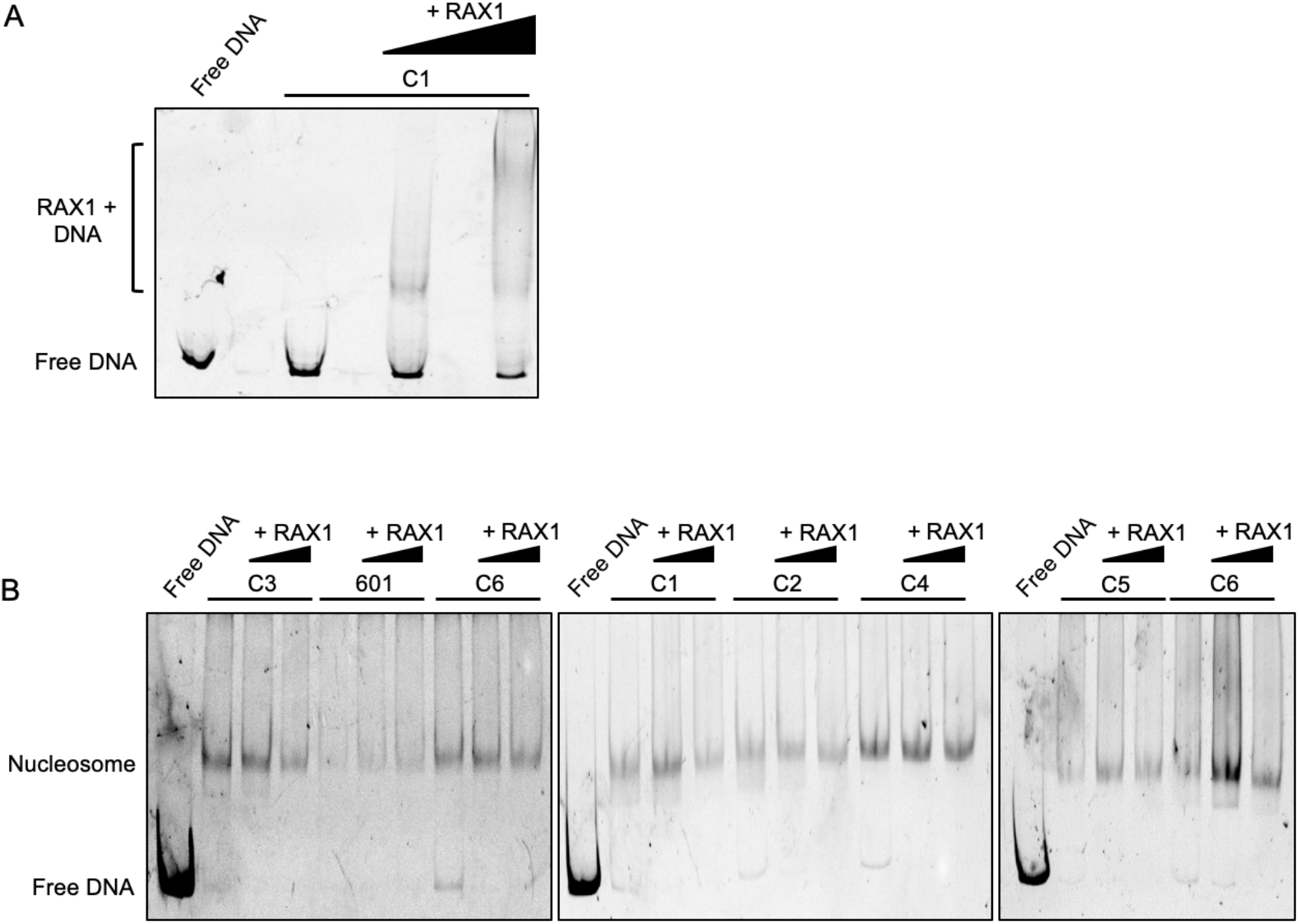
EMSA testing RAX1 nucleosomal DNA binding on 601 sequence with a RAX binding site inserted at different positions. (A) EMSA with different concentrations of RAX1 (250 nM and 1 μM) and free 601 DNA with a RAX1 binding site showing that recombinant RAX1 protein is active. (B) EMSA with RAX1 (250 nM and 1 μM) and 601 nucleosomes without or with a RAX1 binding site inserted at different positions), showing that RAX1 has no nucleosomal DNA binding activity. RAX1 binding site insertion in 601 sequences follows the same principle as for LFYBS illustrated in Figure 3B. DNA sequences are given in Table S3.

**Supplementary Figure 11:**
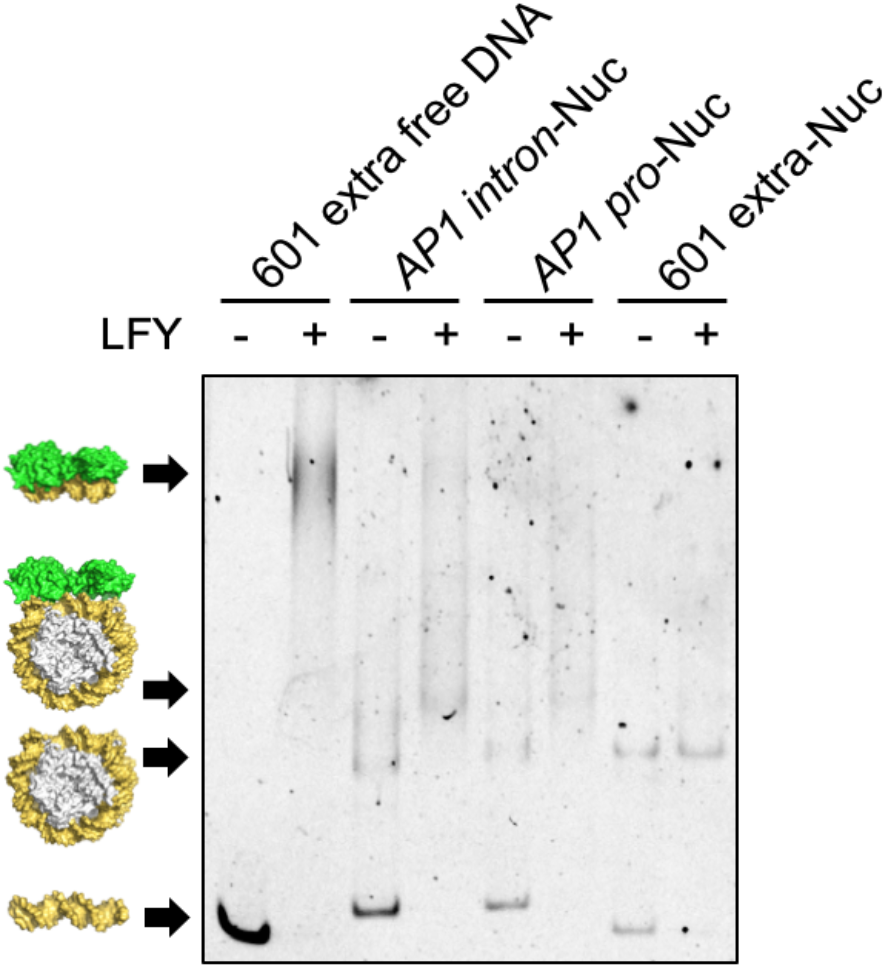
LFY binds nucleosomes assembled with *AP1* intron and promoter sequences (related to Figure 3E). Free DNA or reconstructed nucleosomes are indicated on top of the lanes. The mention extra indicates that the sequence used is longer due to the presence of the amplification primers used for *AP1* sequences (see table 2). The last two lanes are controls showing that these extra sequences do not bind to LFY. Arrows and drawings on the side indicate from bottom to up are free DNA, nucleosome alone, LFY nucleosome complex, and free DNA LFY complex.

**Supplementary Table 1.**
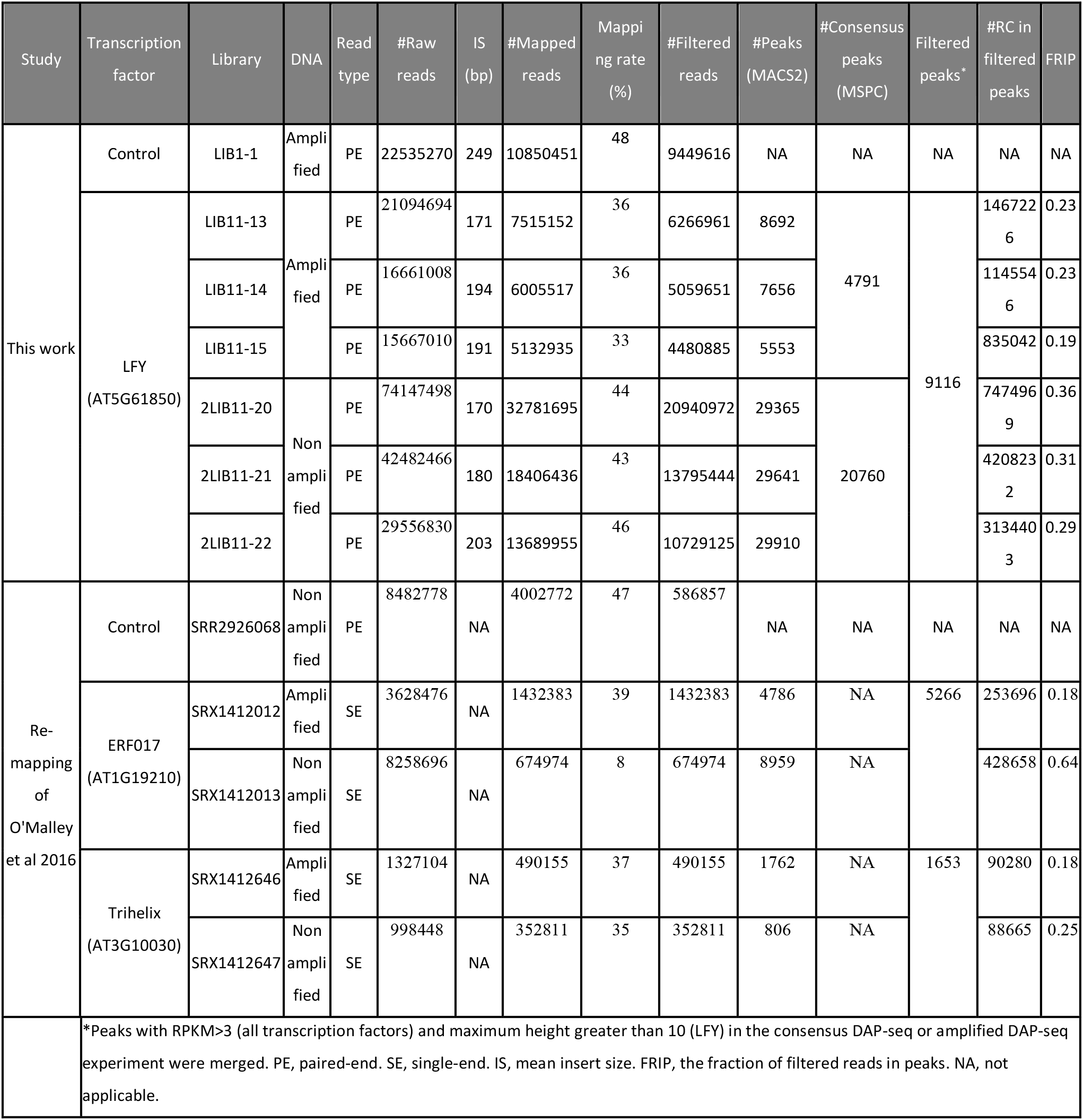
Mapping and peak calling statistics of DAP-seq and ampDAP-seq experiments

**Supplementary Table 2.**
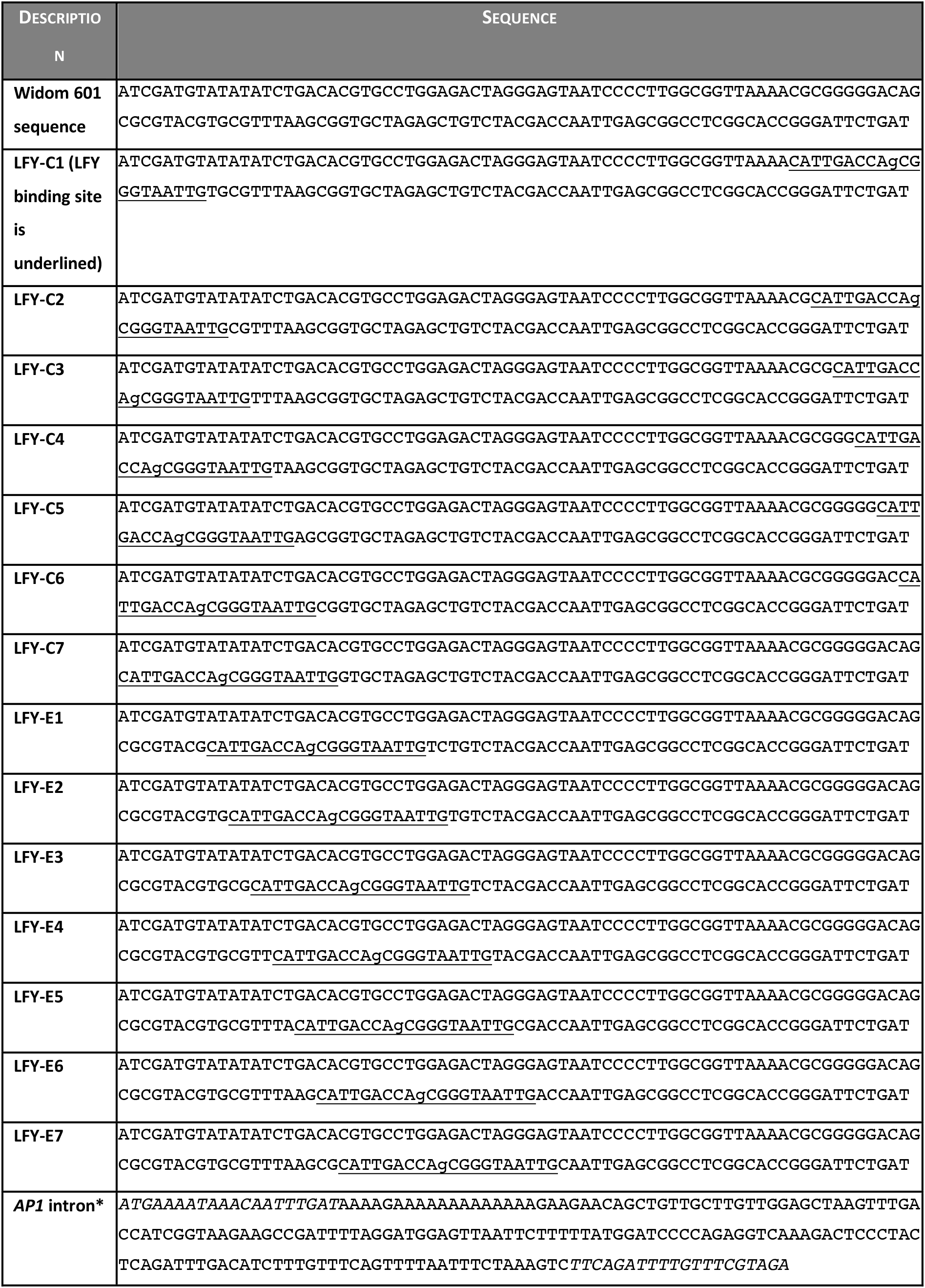
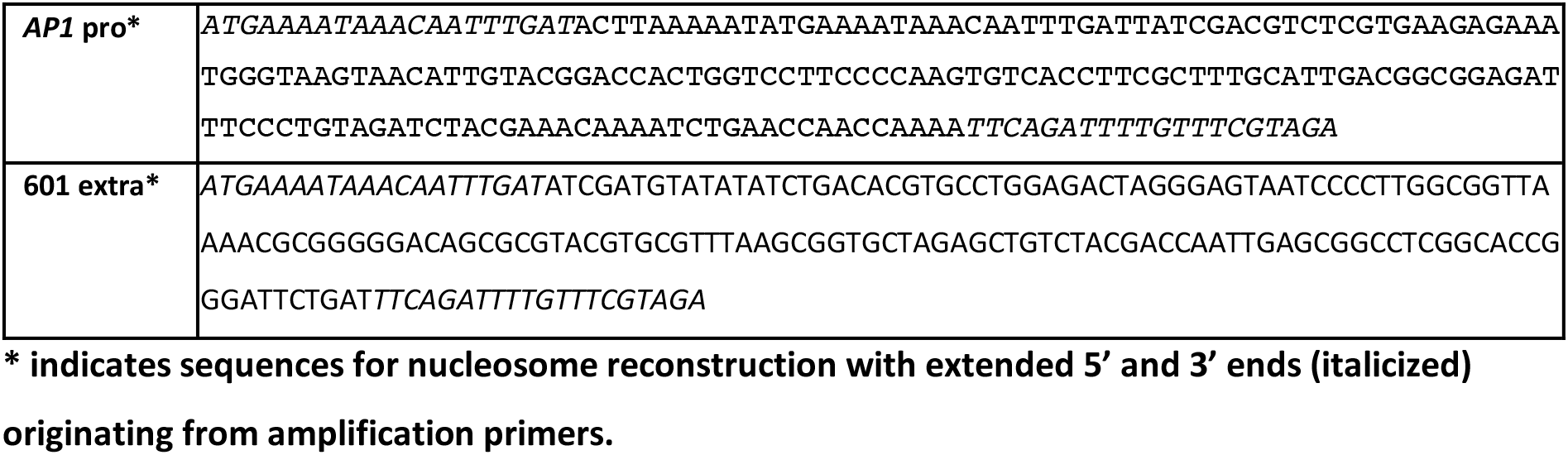
DNA sequences used for nucleosome reconstruction with LFY binding site underlined

**Supplementary Table 3.**
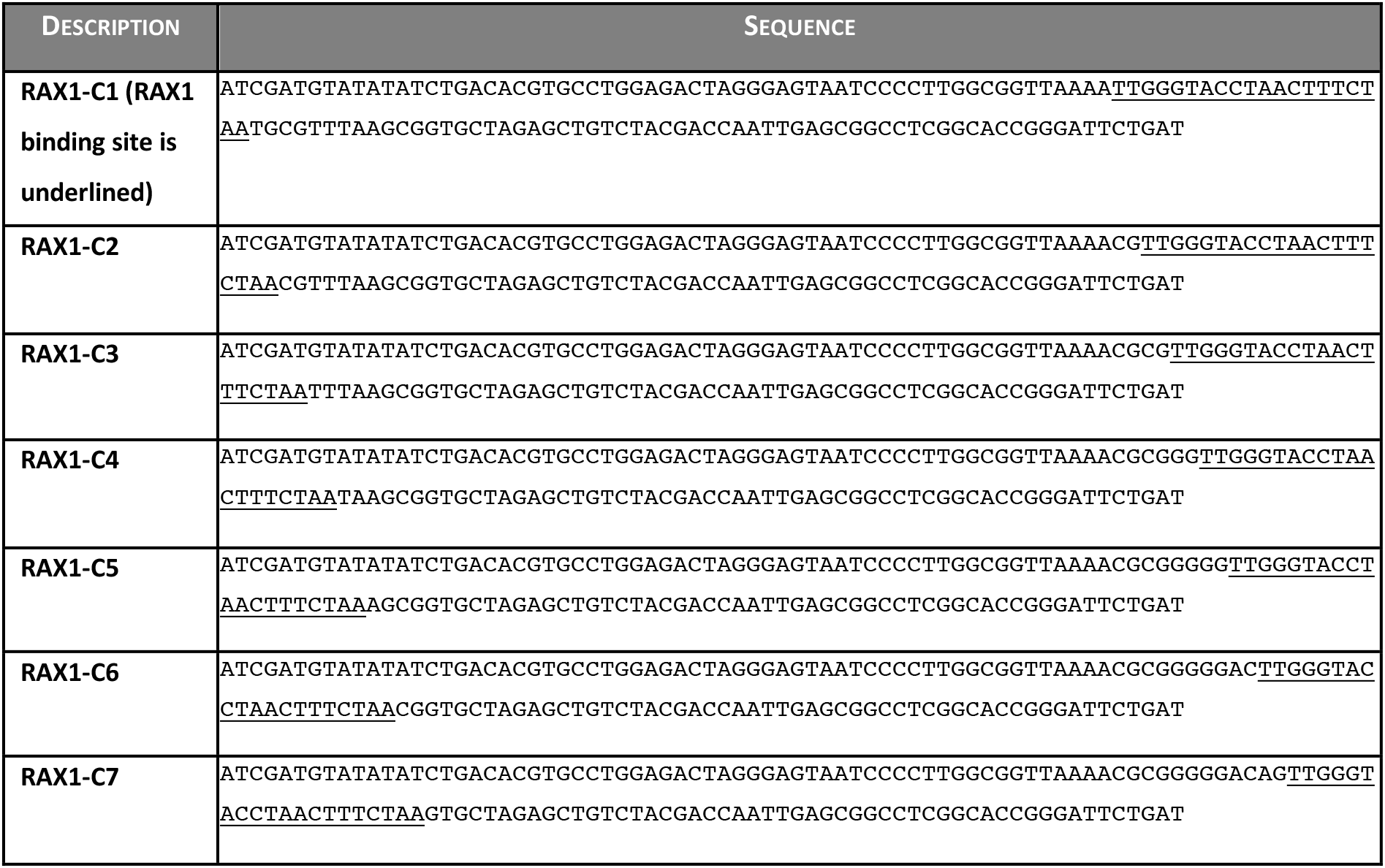
DNA sequences used for nucleosome reconstruction with RAX1 binding site underlined

## References

Archacki, R., Yatusevich, R., Buszewicz, D., Krzyczmonik, K., Patryn, J., Iwanicka-Nowicka, R., Biecek, P., Wilczynski, B., Koblowska, M., Jerzmanowski, A., et al. (2017). Arabidopsis SWI/SNF chromatin remodeling complex binds both promoters and terminators to regulate gene expression. Nuc. Acids Res. 45(6):3116–3129.

Bartlett, A., O’Malley, R. C., Huang, S. C., Galli, M., Nery, J. R., Gallavotti, A., and Ecker, J. R. (2017). Mapping genome-wide transcription-factor binding sites using DAP-seq. Nat. Protocols 12:1659–1672.

Benlloch, R., Kim, M. C., Sayou, C., Thévenon, E., Parcy, F., and Nilsson, O. (2011). Integrating long-day flowering signals: A LEAFY binding site is essential for proper photoperiodic activation of APETALA1. Plant J. 67:1094–1102.

Calonje, M., Sanchez, R., Chen, L., and Sung, Z. R. (2008). EMBRYONIC FLOWER1 Participates in Polycomb Group–Mediated *AG* Gene Silencing in *Arabidopsis*. Plant Cell 20:277–291.

Chae, E., Tan, Q. K.-G., Hill, T. A., and Irish, V. F. (2008). An Arabidopsis F-box protein acts as a transcriptional co-factor to regulate floral development. Development 135:1235–1245.

Chahtane, H., Vachon, G., Le Masson, M., Thévenon, E., Périgon, S., Mihajlovic, N., Kalinina, A., Michard, R., Moyroud, E., Monniaux, M., et al. (2013). A variant of LEAFY reveals its capacity to stimulate meristem development by inducing RAX1. Plant J. 74:678–689.

Chodavarapu, R. K., Feng, S., Bernatavichute, Y. V., Chen, P.-Y., Stroud, H., Yu, Y., Hetzel, J. A., Kuo, F., Kim, J., Cokus, S. J., et al. (2010). Relationship between nucleosome positioning and DNA methylation. Nature 466:388–392.

Chua, E. Y. D., Vasudevan, D., Davey, G. E., Wu, B., and Davey, C. A. (2012). The mechanics behind DNA sequence-dependent properties of the nucleosome. Nuc. Acids Res. 40:6338–6352.

Denay, G., Chahtane, H., Tichtinsky, G., and Parcy, F. (2017). A flower is born: an update on Arabidopsis floral meristem formation. Cur. Op. Plant Biol. 35:15–22.

Dodonova, S. O., Zhu, F., Dienemann, C., Taipale, J., and Cramer, P. (2020). Nucleosome-bound SOX2 and SOX11 structures elucidate pioneer factor function. Nature. 580, 669–672

Fernandez Garcia, M., Moore, C. D., Schulz, K. N., Alberto, O., Donague, G., Harrison, M. M., Zhu, H., and Zaret, K. S. (2019). Structural Features of Transcription Factors Associating with Nucleosome Binding. Mol. Cell 75, 5, 921–932.

Gallois, J.-L., Nora, F. R., Mizukami, Y., and Sablowski, R. (2004). WUSCHEL induces shoot stem cell activity and developmental plasticity in the root meristem. Genes & Dev. 18:375–380.

Gaspar, J. M. (2018). NGmerge: merging paired-end reads via novel empirically-derived models of sequencing errors. BMC Bioinformatics 19:536.

Goodrich, J., Puangsomlee, P., Martin, M., Long, D., Meyerowitz, E. M., and Coupland, G. (1997). A Polycomb-group gene regulates homeotic gene expression in Arabidopsis. Nature 386:44–51.

Goslin, K., Zheng, B., Serrano-Mislata, A., Rae, L., Ryan, P. T., Kwaśniewska, K., Thomson, B., Ó’Maoiléidigh, D. S., Madueño, F., Wellmer, F., et al. (2017). Transcription Factor Interplay between LEAFY and APETALA1/CAULIFLOWER during Floral Initiation. Plant Physiol. 174:1097–1109.

Hamès, C., Ptchelkine, D., Grimm, C., Thevenon, E., Moyroud, E., Gérard, F., Martiel, J.-L., Benlloch, R., Parcy, F., and Müller, C. W. (2008). Structural basis for LEAFY floral switch function and similarity with helix-turn-helix proteins. EMBO J. 27:2628–2637.

Hu, G., Schones, D. E., Cui, K., Ybarra, R., Northrup, D., Tang, Q., Gattinoni, L., Restifo, N. P., Huang, S., and Zhao, K. (2011). Regulation of nucleosome landscape and transcription factor targeting at tissue-specific enhancers by BRG1. Genome Res. 21:1650–1658.

Hu, S., Wan, J., Su, Y., Song, Q., Zeng, Y., Nguyen, H. N., Shin, J., Cox, E., Rho, H. S., Woodard, C., et al. (2013). DNA methylation presents distinct binding sites for human transcription factors. eLife 2:e00726.

Iwafuchi-Doi, M. (2018). The mechanistic basis for chromatin regulation by pioneer transcription factors. Wiley Interdisciplinary Reviews: Systems Biology and Medicine 11(1):e1427.

Iwafuchi-Doi, M., and Zaret, K. S. (2014). Pioneer transcription factors in cell reprogramming. Genes Dev. 28:989–998.

Iwafuchi-Doi, M., and Zaret, K. S. (2016). Cell fate control by pioneer transcription factors. Development 143:1833–1837.

Jalili, V., Matteucci, M., Masseroli, M., and Morelli, M. J. (2015). Using combined evidence from replicates to evaluate ChIP-seq peaks. Bioinformatics 31:2761–2769.

Jin, R., Klasfeld, S., Garcia, M. F., Xiao, J., Han, S.-K., Konkol, A., Zhu, Y., and Wagner, D. (2020). LEAFY is a pioneer transcription factor and licenses cell reprogramming to floral fate. BioRxiv Advance Access published March 18, 2020, doi:10.1101/2020.03.16.994418.

Kaufmann, K., Pajoro, A., and Angenent, G. C. (2010). Regulation of transcription in plants: mechanisms controlling developmental switches. Nat. Rev. Genet. 11:830–842.

King, H. W., and Klose, R. J. (2017). The pioneer factor OCT4 requires the chromatin remodeller BRG1 to support gene regulatory element function in mouse embryonic stem cells. eLife 6:1–24.

Klemm, S. L., Shipony, Z., and Greenleaf, W. J. (2019). Chromatin accessibility and the regulatory epigenome. Nat. Rev. Genet. 20:207–220.

Lai, X., Stigliani, A., Lucas, J., Hugouvieux, V., Parcy, F., and Zubieta, C. (2020). Genome-wide binding of SEPALLATA3 and AGAMOUS complexes determined by sequential DNA-affinity purification sequencing. Nuc. Acids Res. 48:9637–9648.

Langmead, B., and Salzberg, S. L. (2012). Fast gapped-read alignment with Bowtie 2. Nature Methods 9:357–359.

Levin, J.Z. and Meyerowitz, E. M. (1995). UFO: An Arabidopsis Gene lnvolved in Both Floral Meristem and Floral Organ Development. Plant Cell 7:529–548.

Li, H., Handsaker, B., Wysoker, A., Fennell, T., Ruan, J., Homer, N., Marth, G., Abecasis, G., Durbin, R., and 1000 Genome Project Data Processing Subgroup (2009). The Sequence Alignment/Map format and SAMtools. Bioinformatics 25:2078–2079.

Li, S., Bo Zheng, E., Zhao, L., and Liu, S. (2019). Nonreciprocal and Conditional Cooperativity Directs the Pioneer Activity of Pluripotency Transcription Factors. Cell Reports 28, 10: 2689–2703.e4

Liu, Y., Olanrewaju, Y. O., Zheng, Y., Hashimoto, H., Blumenthal, R. M., Zhang, X., and Cheng, X. (2014). Structural basis for Klf4 recognition of methylated DNA. Nuc. Acids Res. 42:4859–4867.

Lohmann, J. U., Hong, R. L., Hobe, M., Busch, M. A., Parcy, F., Simon, R., and Weigel, D. (2001). A Molecular Link between Stem Cell Regulation and Floral Patterning in Arabidopsis. Cell 105:793–803.

Lowary, P. T., and Widom, J. (1998). New DNA sequence rules for high affinity binding to histone octamer and sequence-directed nucleosome positioning. J. Mol. Biol. 276:19–42.

Machanick, P., and Bailey, T. L. (2011). MEME-ChIP: Motif analysis of large DNA datasets. Bioinformatics 27:1696–1697.

Magnani, L., Eeckhoute, J., and Lupien, M. (2011). Pioneer factors: Directing transcriptional regulators within the chromatin environment. Trends Genet. 27:465–474.

Mayran, A., and Drouin, J. (2018). Pioneer transcription factors shape the epigenetic landscape. J. Biol. Chem. 7293(36):13795–13804.

Mayran, A., Khetchoumian, K., Hariri, F., Pastinen, T., Gauthier, Y., Balsalobre, A., and Drouin, J. (2018). Pioneer factor Pax7 deploys a stable enhancer repertoire for specification of cell fate. Nat. Genet. 50: 259–269.

McGinty, R. K., and Tan, S. (2015). Nucleosome Structure and Function. Chem. Rev. 115:2255–2273.

Michael, A. K., Grand, R. S., Isbel, L., Cavadini, S., Kozicka, Z., Kempf, G., Bunker, R. D., Schenk, A. D., Graff-Meyer, A., Pathare, G. R., et al. (2020). Mechanisms of OCT4-SOX2 motif readout on nucleosomes. Science 368, 6498:1460–1465.

Moreau, F., Thévenon, E., Blanvillain, R., Lopez-Vidriero, I., Franco-Zorrilla, J. M., Dumas, R., Parcy, F., Morel, P., Trehin, C., and Carles, C. C. (2016). The Myb-domain protein ULTRAPETALA1 INTERACTING FACTOR 1 controls floral meristem activities in *Arabidopsis*. Development 143:1108–1119.

Moyroud, E., Kusters, E., Monniaux, M., Koes, R., and Parcy, F. (2010). LEAFY blossoms. Trends Plant Sci. 15:346–352.

Moyroud, E., Minguet, E. G., Ott, F., Yant, L., Posé, D., Monniaux, M., Blanchet, S., Bastien, et al. (2011). Prediction of regulatory interactions from genome sequences using a biophysical model for the Arabidopsis LEAFY transcription factor. Plant Cell 23:1293–1306.

Okuwaki, M., Kato, K., Shimahara, H., Tate, S., and Nagata, K. (2005). Assembly and Disassembly of Nucleosome Core Particles Containing Histone Variants by Human Nucleosome Assembly Protein I. MCB 25:10639–10651.

O’Malley, R. C., Huang, S. shan C., Song, L., Lewsey, M. G., Bartlett, A., Nery, J. R., Galli, M., Gallavotti, A., and Ecker, J. R. (2016). Cistrome and Epicistrome Features Shape the Regulatory DNA Landscape. Cell 166:1598.

Pajoro, A., Madrigal, P., Muiño, J. M., Matus, J. T., Jin, J., Mecchia, M. A., Debernardi, J. M., Palatnik, J. F., Balazadeh, S., Arif, M., et al. (2014). Dynamics of chromatin accessibility and gene regulation by MADS-domain transcription factors in flower development. Genome Biol. 15:R41.

Parcy, F., Nilsson, O., Busch, M. A., Lee, I., and Weigel, D. (1998). A genetic framework for floral patterning. Nature 395:561–566.

Quinlan, A. R., and Hall, I. M. (2010). BEDTools: a flexible suite of utilities for comparing genomic features. Bioinformatics 26:841–842.

Risseeuw, E., Venglat, P., Xiang, D., Komendant, K., Daskalchuk, T., Babic, V., Crosby, W., and Datla, R. (2013). An activated form of UFO alters leaf development and produces ectopic floral and inflorescence meristems. PLoS ONE 8.

Sayou, C., Nanao, M. H., Jamin, M., Pose, D., Thevenon, E., Gregoire, L., Tichtinsky, G., Denay, G., Ott, F., Llobet, M. P., et al. (2016). A SAM oligomerization domain shapes the genomic binding landscape of the LEAFY transcription factor. Nat. Commun. 7:11222.

Sequeira-Mendes, J., Araguez, I., Peiro, R., Mendez-Giraldez, R., Zhang, X., Jacobsen, S. E., Bastolla, U., and Gutierrez, C. (2014). The Functional Topography of the Arabidopsis Genome Is Organized in a Reduced Number of Linear Motifs of Chromatin States. Plant Cell 26:2351–2366.

Sherwood, R. I., Hashimoto, T., O’Donnell, C. W., Lewis, S., Barkal, A. A., Van Hoff, J. P., Karun, V., Jaakkola, T., and Gifford, D. K. (2014). Discovery of directional and nondirectional pioneer transcription factors by modeling DNase profile magnitude and shape. Nat. Biotech. 32:171–178.

Shim, Y., Duan, M.-R., Chen, X., Smerdon, M. J., and Min, J.-H. (2012). Polycistronic coexpression and nondenaturing purification of histone octamers. Ana. Biochem. 427:190–192.

Slattery, M., Zhou, T., Yang, L., Dantas Machado, A. C., Gordân, R., and Rohs, R. (2014). Absence of a simple code: How transcription factors read the genome. Trends Biochem. Sci. 39:381–399.

Soufi, A., Garcia, M. F., Jaroszewicz, A., Osman, N., Pellegrini, M., and Zaret, K. S. (2015). Pioneer transcription factors target partial DNA motifs on nucleosomes to initiate reprogramming. Cell 161:555–568.

Tao, Z., Shen, L., Gu, X., Wang, Y., Yu, H., and He, Y. (2017). Embryonic epigenetic reprogramming by a pioneer transcription factor in plants. Nature 551:124–128.

Turck, F., Roudier, F., Farrona, Sara., Martin-Magniette, M.-L., Guillaume, E., Buisine, N., Gagnot, S., Martienssen, R. A., Coupland, G., and Colot, V. (2007). Arabidopsis TFL2/LHP1 Specifically Associates with Genes Marked by Trimethylation of Histone H3 Lysine 27. PLoS Genetics 3(6):e86.

Wagner, D. (1999). Transcriptional Activation of APETALA1 by LEAFY. Science 285:582–584.

Weber, E., Buzovetsky, O., Heston, L., Yu, K.-P., Knecht, K. M., El-Guindy, A., Miller, G., and Xiong, Y. (2019). A Noncanonical Basic Motif of Epstein-Barr Virus ZEBRA Protein Facilitates Recognition of Methylated DNA, High-Affinity DNA Binding, and Lytic Activation. J Virol 93:e00724–19.

Winter, C. M., Austin, R. S., Blanvillain-Baufumé, S., Reback, M. A., Monniaux, M., Wu, M. F., Sang, Y., Yamaguchi, A., Yamaguchi, N., Parker, J. E., et al. (2011). LEAFY Target Genes Reveal Floral Regulatory Logic, cis Motifs, and a Link to Biotic Stimulus Response. Dev. Cell 20:430–443.

Wu, M., Sang, Y., Bezhani, S., Yamaguchi, N., Han, S., Li, Z., Su, Y., Slewinski, T. L., and Wagner, D. (2012). SWI2/SNF2 chromatin remodeling ATPases overcome polycomb repression and control floral organ identity with the LEAFY and SEPALLATA3 transcription factors. Proc. Nat. Acad. Sci. USA 109:3576–3581.

Yamaguchi, N., Wu, M. F., Winter, C. M., Berns, M. C., Nole-Wilson, S., Yamaguchi, A., Coupland, G., Krizek, B. A., and Wagner, D. (2013). A Molecular Framework for Auxin-Mediated Initiation of Flower Primordia. Dev. Cell 24:271–282.

Yin, Y., Morgunova, E., Jolma, A., Kaasinen, E., Sahu, B., Khund-Sayeed, S., Das, P. K., Kivioja, T., Dave, K., Zhong, F., et al. (2017). Impact of cytosine methylation on DNA binding specificities of human transcription factors. Science 356:eaaj2239.

Zaret, K. S. (2020). Pioneer Transcription Factors Initiating Gene Network Changes. Annu. Rev. Genet. 54:annurev-genet-030220-015007.

Zhang, Y., Liu, T., Meyer, C. A., Eeckhoute, J., Johnson, D. S., Bernstein, B. E., Nussbaum, C., Myers, R. M., Brown, M., Li, W., et al. (2008). Model-based Analysis of ChIP-Seq (MACS). Genome Biol. 9:R137.

Zhang, W., Zhang, T., Wu, Y., and Jiang, J. (2012). Genome-Wide Identification of Regulatory DNA Elements and Protein-Binding Footprints Using Signatures of Open Chromatin in Arabidopsis. Plant Cell 24:2719–2731.

Zhang, T., Zhang, W., and Jiang, J. (2015). Genome-Wide Nucleosome Occupancy and Positioning and Their Impact on Gene Expression and Evolution in Plants. Plant Physiol. 168:1406–1416.

Zhang, Q., Wang, D., Lang, Z., He, L., Yang, L., Zeng, L., Li, Y., Zhao, C., Huang, H., Zhang, H., et al. (2016). Methylation interactions in Arabidopsis hybrids require RNA-directed DNA methylation and are influenced by genetic variation. Proc. Natl. Acad. Sci. USA 113:E4248–E4256.

Zhu, H., Wang, G., and Qian, J. (2016). Transcription factors as readers and effectors of DNA methylation. Nat. Rev. Genet. 17:551–565.

